# Single cell RNA-seq by mostly-natural sequencing by synthesis

**DOI:** 10.1101/2022.05.29.493705

**Authors:** Sean K. Simmons, Gila Lithwick-Yanai, Xian Adiconis, Florian Oberstrass, Nika Iremadze, Kathryn Geiger-Schuller, Pratiksha I. Thakore, Chris J. Frangieh, Omer Barad, Gilad Almogy, Orit Rozenblatt-Rosen, Aviv Regev, Doron Lipson, Joshua Z. Levin

## Abstract

Massively parallel single cell RNA-seq (scRNA-seq) for diverse applications, from cell atlases to functional screens, is increasingly limited by sequencing costs, and large-scale low-cost sequencing can open many additional applications, including patient diagnostics and drug screens. Here, we adapted and systematically benchmarked a newly developed, mostly-natural sequencing by synthesis method for scRNA-seq. We demonstrate successful application in four scRNA-seq case studies of different technical and biological types, including 5’ and 3’ scRNA-seq, human peripheral blood mononuclear cells from a single individual and in multiplex, as well as Perturb-Seq. Our data show comparable results to existing technology, including compatibility with state-of-the-art scRNA-seq libraries independent of the sequencing technology used – thus providing an enhanced cost-effective path for large scale scRNA-seq.

---

Cellular profiling by single-cell RNA-seq (scRNA-seq) now enables studying and characterizing cellular states and pathways at ever-growing experimental scales, including the Human Cell Atlas ^1^, cell atlases for tumors ^2^ and other diseases ^3, 4^, and large-scale Perturb-seq screens of millions of cells under genetic ^5, 6^ or drug ^7^ perturbations. As methods for capturing and processing single cell libraries have been radically scaled in the past few years ^8-11^, sequencing technologies are becoming a major barrier to the broad adoption of scRNA-seq in both basic research and the clinic.

Here, we describe the application of a new sequencing technology that has the potential to advance single cell genomics by significantly lowering the sequencing cost component of scRNA-seq. Mostly-natural sequencing by synthesis (mnSBS) is a new sequencing chemistry that leverages a low fraction of labeled nucleotides to combine the efficiency of non-terminating chemistry with the throughput and scalability of optical endpoint scanning to enable low-cost, high-throughput sequencing. To benchmark mnSBS with scRNA-seq, we performed experiments with four library types, sequenced in parallel on an Illumina sequencer and on an Ultima Genomics (Ultima) prototype sequencer implementing mnSBS (**Fig. 1a**).

**Fig. 1.**
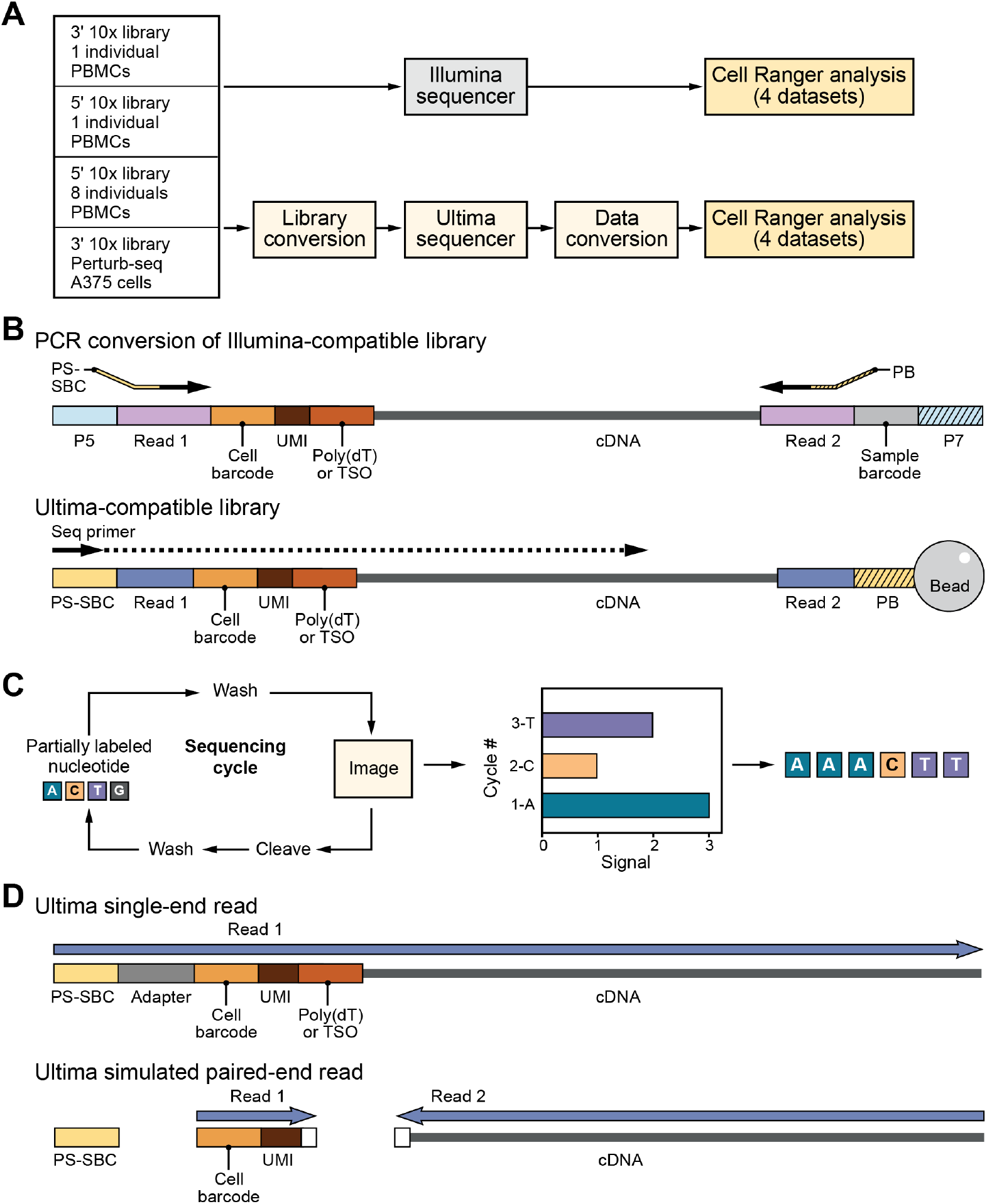
Experimental Design. (**a**) Workflow showing 4 samples used and adjustments made for Ultima sequencing. (**b**) Library conversion showing PCR process to change adapters from Illumina (P5 and P7, parts of Read 1 and 2) to Ultima (PS-SBC and PB, parts of Read 1 and 2). 5’ libraries have Template Switch Oligo (TSO) and 3’ libraries have poly(dT). (**c**) Mostly-natural sequencing by synthesis schematic. (**d**) Data conversion of single-end reads to simulated paired-end reads needed for Cell Ranger analysis. White box shows 5 bases trimmed from cDNA and 3 bases trimmed from UMI adjacent to the poly(dT) sequence in 3’ libraries. In 5’ libraries, only 3 bases were trimmed from the cDNA next to the TSO. PS-SBC read is used to deconvolute multiplexed libraries.

To implement mnSBS for massively-parallel, droplet-based scRNA-seq, we converted a typical scRNA-seq workflow to be compatible with Ultima sequencing (**Fig. 1b**-**d**; **Online Methods**). Focusing on 10x Chromium scRNA-seq (**Online Methods**), a popular method, we first added adapters to cDNA libraries specific for Ultima sequencing (**Fig. 1b**). Next, we address the fact that droplet-based scRNA-seq relies on pairing each cDNA read with a cell barcode (CBC) and a Unique Molecular Identifier (UMI) (**Online Methods**). With Illumina sequencing, the two ends of the library are sequenced separately by paired-end sequencing, but for single-end Ultima sequencing, we capture all the information in a single 200 to 250 base read (**Fig. 1d**), such that the CBC and UMI are read first and followed by the cDNA. For those reads derived from the transcript’s 3’ end, we sequence through poly(T) bases, which are due to the mRNA poly(A) tail, adjacent to the cDNA sequence.

To evaluate mnSBS with scRNA-seq, we carried out experiments with four libraries, spanning different technical and biological use cases, and sequenced each in parallel on both Ultima and Illumina sequencers (**Online Methods**). Three libraries were from peripheral blood mononuclear cells (PBMCs) of healthy human donors, spanning 3’ scRNA-seq (∼7,000 cells, 1 individual), 5’ scRNA-Seq (∼7,000 cells, 1 individual), and a library generated in multiplex by pooling cells from eight donors (∼24,000 cells, 8 individuals, 5’ scRNA-seq). We chose PBMCs because they are primary human cells, include diverse cell types of various sizes and frequencies, and have been used for previous benchmarking ^12, 13^. The fourth library was from a Perturb-Seq ^5, 6^ experiment, where ∼20,000 cells were profiled after CRISPR/Cas9 pooled genetic perturbation, followed by scRNA-Seq to detect both the cell’s profile and associated guide RNA. Together, the four libraries span three major use cases – individual patient atlas, multiplex patient profiling, and large-scale screens, and the two most commonly used library types for scRNA-seq.

We first tested the feasibility of mnSBS for scRNA-seq, with matched Ultima and 5’ and 3’ droplet-based scRNA-seq of PBMCs. Initial analysis (**Online Methods**) showed that the number of UMIs generated at a given sequencing depth was comparable between Ultima and Illumina in the 5’ libraries, while for the 3’ libraries we obtained more UMIs with Illumina than Ultima (**Fig. 2a**), due to sequence quality differences. While Ultima and Illumina data for 5’ libraries were similar, for the 3’ data there was lower quality for Ultima in the bases flanking the poly(T) region – the 3’ end of the UMI and the 5’ end of the cDNA (**Extended Data Fig. 1a**). Indeed, filtering out reads that have bases with quality <10 in their UMI (the filter applied by the pre-processing pipeline we used, Cell Ranger ^14^), yields similar rarefaction curves for Illumina and Ultima (**Extended Data Fig. 1b**). Thus, much of the difference in the observed number of UMIs per sequenced read for 3’ libraries is explained by the lower sequence quality UMIs in the Ultima data due to the need to sequence through the poly(T) bases. To overcome this, for 3’ libraries we trimmed 5 bases from the cDNA adjacent to the poly(T) bases and then explored how best to trim the UMI. As we shortened the UMIs, UMIs that differed only in the trimmed bases “collapsed” into a single UMI leading to decreases in the fraction of UMI/CBC pairs that occur in only one gene at roughly the same rate in Illumina and Ultima data (**Extended Data Fig. 1c**). Shortening the UMIs for Illumina had a minimal effect at 9 bases or more (**Extended Data Fig. 1d**), suggesting that the challenges with Ultima reads were due to lower base quality and that trimming to 9 bases was reasonable. This led us to exclude the last 3 bases of each UMI in Ultima 3’ data in subsequent downstream analysis (**Online Methods**).

**Fig. 2.**
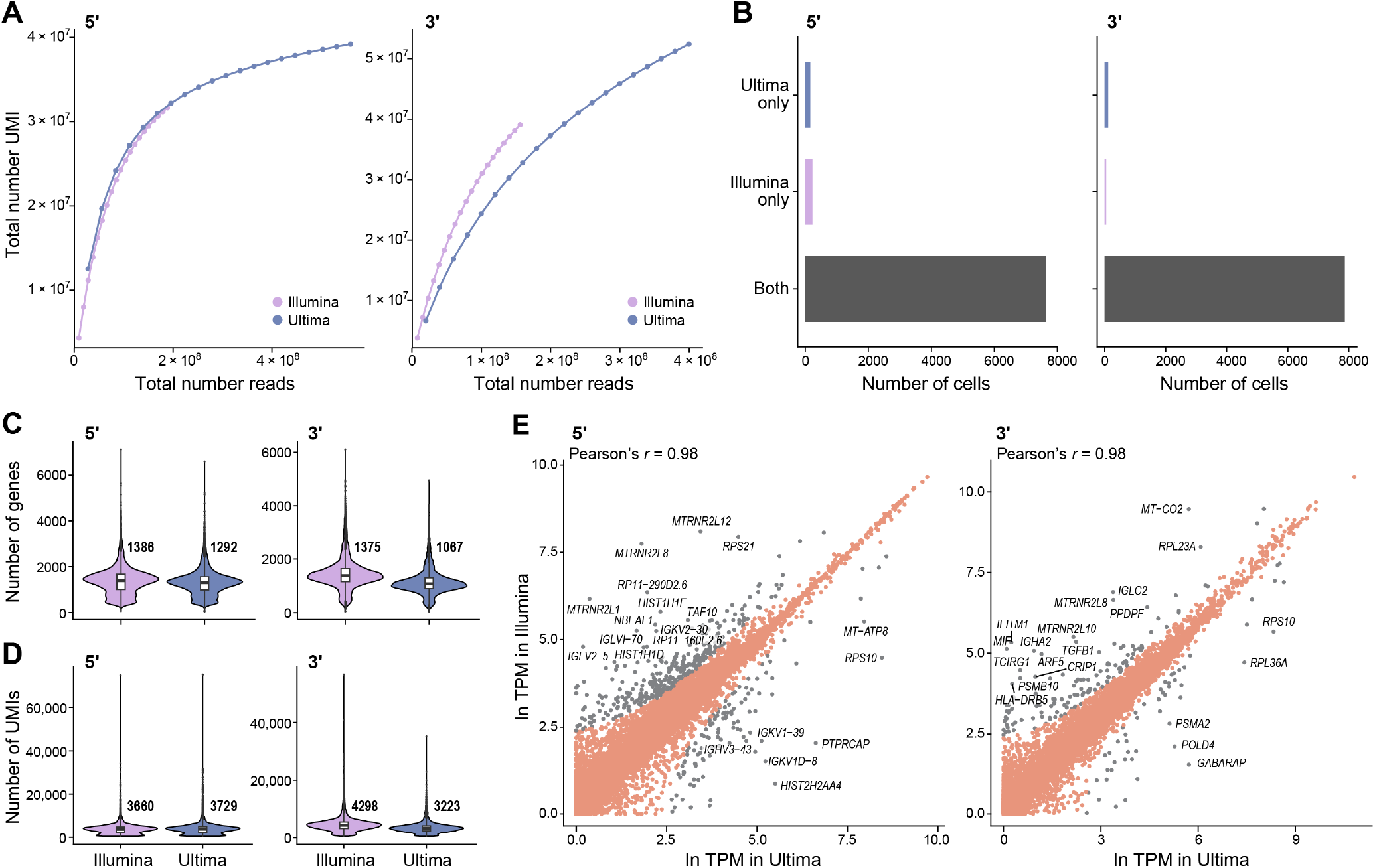
Quality Metrics for matched 5’ and 3’ libraries. (**a**) Total number of UMIs detected per cell at different sequencing depths. For (**b**) – (**e**) reads were sampled so that Illumina and Ultima have the same number of reads. (**b**) Number of cells identified by Cell Ranger only in Ultima, only in Illumina, or both. Distribution of the number of genes (**c**) or UMIs (**d**) per cell. Boxplots defined by 25% and 75% quantiles with the median marked in between. (**e**) Scatter plots with one point for each gene. Labeled genes (grey) have a high fold change (FC) (FC >2 using a pseudocount of 10 TPM). The 20 genes with the highest FC are labeled in each plot. For all 3’ libraries, the last three UMI bases were trimmed for quality reasons.

Next, comparing the performance of these PBMC 3’ and 5’ matched libraries, we obtained similar overall performance for both sequencing technologies. First, to correct for differences in sequencing depths, which were higher in Ultima than Illumina, we randomly sampled Ultima reads, so that we used the same number of reads for each sequencing platform (**Online Methods**). Both technologies identified nearly all the same CBCs (**Fig. 2b**, 7,916 cells (Ultima) vs 7,926 cells (Illumina) in the 3’ data, and 7,875 cells (Ultima) vs 7,854 cells (Illumina) in the 5’ data), with the same number of UMIs and genes per cell for 5’ libraries and slightly lower numbers for 3’ libraries with Ultima (as expected) (**Fig. 2c,d**). When we sampled reads to have the same number of UMIs (**Online Methods**), we obtained a similar number of genes per cell in Illumina and Ultima also for 3’ libraries (**Extended Data Fig. 2**). Other metrics (**Supplementary Table 1**) also showed similar overall performance, with slightly higher genome mapping rates in Ultima but comparable transcriptome mapping rates.

The two technologies yielded highly correlated expression levels, albeit with some outlier genes and minor differences (Pearson’s *r*=0.98 in all cases; **Fig. 2e, Extended Data Fig. 2c**). (As expected, when a single sequencing run was randomly split into two datasets, we see even higher correlation of expression levels (**Extended Data Fig. 2d**)). Specifically, there was a modest bias, particularly in the 3’ libraries, towards genes with higher GC content having higher expression in Illumina and the longest genes having higher expression in Ultima 3’ libraries (**Extended Data Fig. 3a,b**). Of the 166 genes with differences in expression for 3’ PBMC between the two sequencing platforms, most (130 genes, 78.3%) differed in the fraction of reads that were assigned by Cell Ranger to the gene out of all the reads mapped to that gene region (**Extended Data Fig. 3c**). This is likely related to how Ultima and Illumina reads map to different locations relative to the transcript, as expected from the difference in single-end *vs*. paired-end reads (**Fig. 1d**): in 5’ data, Ultima reads map closer to the 5’ end than Illumina reads, while in 3’ data, Ultima reads map closer to the 3’ end than Illumina reads (**Extended Data Fig. 3d,e**). Because Cell Ranger excludes reads that do not fully map within annotated gene boundaries, more Ultima reads are excluded from analysis as they are closer to gene ends (**Extended Data Fig. 3d,e**), as shown for example for *LILRA5* and *HIST1H1D* (**Extended Data Fig. 3f,g**). This difference in location can also lead to more multimapping or ambiguous reads (**Extended Data Fig. 3h, Supplementary Table 2**). For example, four (*ARF5, MIF, IFITM1*, and *TCIRG1*) of the 20 genes with the largest log fold change (logFC) between Ultima and Illumina in the 3’ data (labeled in **Fig 2E**) have higher expression in Illumina and a much higher rate of mapped ambiguous reads in the Ultima than the Illumina data (>50 vs. <10 ambiguous reads per non-ambiguous read for each gene, respectively) (**Supplementary Table 2**), possibly explaining the difference in their expression levels. Shortening Ultima Read 2 to the same length as Illumina Read 2 had a small effect on the fraction of assigned reads (**Extended Data Fig. 3h**) and other metrics (**Supplementary Table 1**) – suggesting read length is not a major factor in the differences we observed.

To further explore the effects of gene annotation on Ultima and Illumina-based scRNA-seq, we extended the standard reference using RNA-seq data, as we have previously shown this can recover the expression of a gene with an alternative 3’ end compared to the annotation ^15^. We created a pipeline that extends the annotated gene boundaries based on reads that overlap a gene but are not completely contained in any of its annotated exons (**Online Methods**). We generated three such references, extended with either (1) published bulk PBMC data ^12^, (2) the Ultima 3’ scRNA-seq data, or (3) the Ultima 5’ scRNA-seq data (with Ultima and Illumina data sampled to the same number of reads). We compared the expression of genes in Ultima data processed with the extended references to those in Illumina data either with or without the extended reference (**Extended Data Fig. 4, Supplementary Tables 1 and 3**). Analyzing the 5’ PBMC data with the extended reference decreased the number of differentially expressed (DE) genes between Ultima and Illumina by 22 to 23% (absolute logFC > ln(2)) compared with the standard reference, while other overall metrics were largely unchanged. In the 3’ data, there were a similar number of DE genes in analyses with the extended and standard references, although the expression of some genes, e.g., *LILRA5* and *MT-CO2*, agreed much more closely using the extended reference. Comparing gene expression levels for the same sequencing dataset processed with the standard or an extended reference shows that most levels are very similar, though a sizeable number (23 to 83) are higher and a few (1 to 3) are lower (**Extended Data Fig. 4c**). Also. some of the top genes that differ between the extended and standard references are genes that differ between Ultima and Illumina with the standard reference, e.g., *MT-CO2* and *LILRA5* in the 3’ data and *HIST1H1D* and *HIST1H1E* in the 5’ data (**Fig. 2e, Extended Data Fig. 3f,g**). This suggests that a data-driven extended reference might help recover expression in Ultima scRNA-seq data, particularly when using 5’ data. Alternatively, one could consider modifying the way Cell Ranger counts UMIs to better take advantage of reads that overlap genes but are not completely contained within them.

To compare the biological insights derived from scRNA-seq using the two technologies, we turned to analyze 5’ scRNA-seq of PBMCs from eight individuals processed together and sequenced with both Ultima and Illumina (**Online Methods**). Both methods have roughly the same number of UMIs in this dataset (<1% difference) and performed similarly (**Supplementary Table 1** and **Extended Data Fig. 5**, using all reads). We also generated matched T cell Receptor (TCR) and B cell Receptor (BCR) Illumina sequencing data (**Online Methods**). Ultima sequencing was not used for this, because the 10x Chromium constructs specifically require paired-ends or much longer single-end reads to cover the entirety of these genes.

In the 8 individuals PBMC dataset, the two sequencing platforms produced very similar results for the common tasks of genotype-based assignment, cell type labeling, and DE gene identification, and were well-embedded together. First, we used Vireo ^16^, which finds genotype clusters in the data without prior knowledge of the genotypes of individuals in the experiment, to assign reads to each individual in the mixture (**Online Methods**). Both Ultima and Illumina data returned highly concordant labels (**Fig. 3a**), with 92% agreement in label if we include those cells declared doublets or unassigned (χ^2^ test for independence gives a p-value < 2.2 × 10^−16^ and χ^2^=199127 with df=81), and >99.9% agreement if the cell is assigned singlet by both technologies (only 5 cells differ, χ^2^ test for independence gives a p-value < 2.2 × 10^−16^ and χ^2^=146879 with df=49). Next, we clustered the cells for each of the two datasets separately (**Online Methods**), and used Azimuth ^17^ to automatically label cell types in each (**Online Methods**). In both sequencing datasets, we identify the major cell types expected for PBMCs, with the expected cell type markers (**Extended Data Fig. 6a,b**), and cells are comparably well-mixed among individuals (**Fig. 3b**), with low adjusted mutual information (AMI) between cell type and individual in both Ultima (0.026) and Illumina (0.025) (AMI = 0 corresponds to no relation between individual and cell type; AMI = 1 corresponds to the case of perfect agreement between the two labelings). The two sequencing datasets also had high agreement on proportions of each cell type from each individual, both for the main cell type categories (**Fig. 3c**, 95% agreement in cell type labels between Ultima and Illumina, χ^2^ test for independence gives a p-value < 2.2 × 10^−16^ and χ^2^=123891 with df=49, AMI = 0.88) and for finer cell subsets, such as subclusters of T cells (**Fig. 3d**). They further agreed on differential expression between cell types (**Fig. 3e**; *r*=0.93-0.95), such that 67.9% of genes that are significantly DE in one cell type in one of the two datasets are significant in both for that cell type. We found similar results with 5’ and 3’ PBMC datasets from a single individual (**Extended Data Fig. 7**). Moreover, the two PBMC mixture datasets were co-embedded well together into a joint 2-dimensional space using Uniform Manifold Approximation and Projection (UMAP) after regressing out dataset of origin (**Online Methods**), with good mixing between datasets (AMI = 0.00068 between the joint clustering and dataset of origin), and good separation of cell types (**Fig. 3f**). Thus, data generated by the two sequencing technologies are compatible and can be combined easily in a single analysis.

**Fig. 3.**
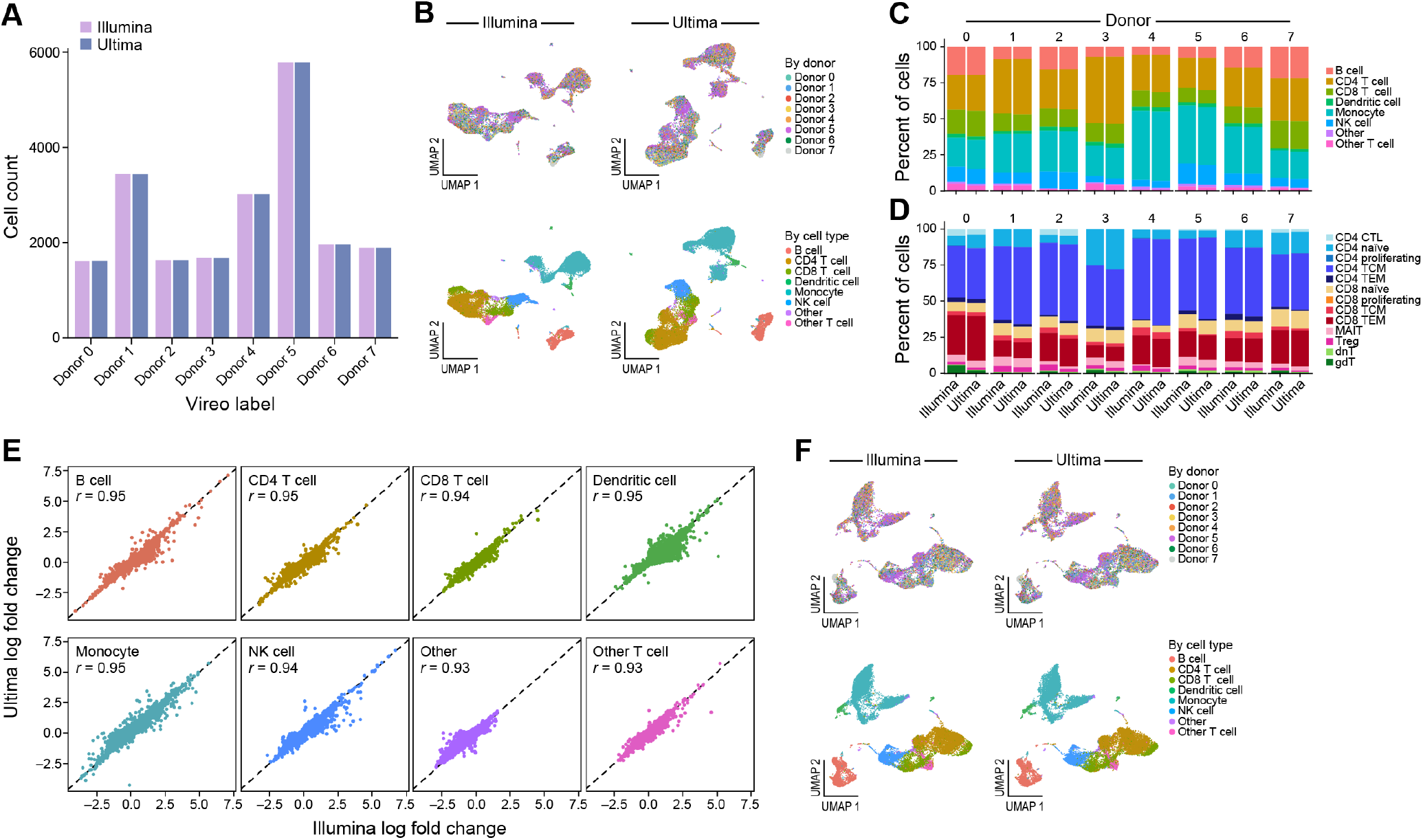
Cell type identification and characterization of a mixture of PBMCs. (**a**) Number of cells assigned to each donor by Vireo. Donors were renamed to match between Ultima and Illumina. (**b**) UMAP plots for Ultima (right) and Illumina (left) colored by donor (top) and cell type (bottom). (**c**) Bar plots of the proportion of each Azimuth-defined cell type in each donor for Ultima and Illumina. (**d**) Bar plots of the proportion of each Azimuth-defined T cell subtype in each donor for Ultima and Illumina. Can see strong agreement. (**e**) Scatter plots of logFC from performing DE between cell type clusters with Presto. (**f**) Joint UMAP of Ultima and Illumina data colored as in (**b**). We did not sample the exact same number of reads from Illumina or Ultima data since they have approximately the same number of total UMIs. NK: Natural Killer cells. CTL: Cytotoxic T cells. TCM: Central memory T cells. TEM: Effector memory T cells. MAIT: Mucosal-associated invariant T cells. Treg: Regulatory T cells. dnT: Double negative T cells. gdT: Gamma delta T cells.

For B and T cells, where we had clonotype assignment by Illumina sequencing of TCRs and BCRs (**Online Methods**), we found good concordance between Ultima and Illumina assignments. Most T cells called by either method had TCR sequences (76% in both Ultima and Illumina) with only a very small percent of cells of other cell types having a TCR sequence (3.7% in Ultima and 3.5% Illumina), with similar results for B cells and BCR sequences (**Extended Data Fig. 6c**; 93% of B cells in both Ultima and Illumina were assigned a BCR clonotype while only 0.72% of non-B cells in Ultima and 0.73% of non-B cells in Illumina were assigned a BCR clonotype). The distribution of T cell subsets to top TCR clonotypes for each individual was also largely concordant between Ultima and Illumina sequencing (**Extended Data Fig. 6d**), with small differences in cell type labeling. CD8 T effector memory (TEM) cells were by far the most likely to be expanded, as expected ^18^. Thus, Ultima sequencing for scRNA-seq can be combined with Illumina sequencing of TCR and BCR genes to generate comparable results to those found with only Illumina sequencing.

To explore finer signals, we compared the two data sets for continuous cell states – such as activation status or the cell cycle – recovered by unsupervised non-negative matrix factorization (NMF). Each NMF factor can reflect a gene program, defined by a non-negative score for each gene (referred to as gene loadings) and a non-negative score for each cell (referred to as cell loadings). Because NMF runs are not identical even when re-run on the same data, to compare NMF models from Ultima and Illumina data, we fit NMF on Ultima data, Illumina data, and a null of randomly permuted Illumina expression values (**Online Methods**), and then measured how well cell or gene loadings fit each dataset. Cell loadings from the model learned on Ultima data fit the Illumina data almost as well as cell loadings from the Illumina-learned model and vice versa, while loadings from the permuted (null) dataset led to much poorer fit (**Extended Data Fig. 8a**). For gene loadings, there was lower performance when fitting data from one sequencing technologies with loadings from a model learned on the data from the other technologies, each to a comparable extent, and both far better than random permutations (**Extended Data Fig. 8b**). Consensus NMF (cNMF) ^19^ (**Online Methods**), which reduces variability due to random sampling between NMF runs, showed high correlations of cell (**Extended Data Fig. 8c**) or gene (**Extended Data Fig. 8d**) loadings between models learned on different runs. The correspondence was comparable to that observed between two independent cNMF runs on the *same* dataset (**Extended Data Fig. 8e,f**), and lower than when comparing a single run to itself (**Extended Data Fig. 8g,h**), as expected. It was also much stronger than comparing cNMF models of two different biological systems (5’ PBMC mixture data and Perturb-seq, see below for details of this experiment) (**Extended Data Fig. 8i,j**). Notably, the same cell subsets score highly for Ultima (**Extended Data Fig. 8k**) and Illumina (**Extended Data Fig. 8l**) data-derived programs on a joint UMAP embedding. For example, factor 13 in Illumina and factor 1 in Ultima scored in the same cells (**Extended Data Fig. 8k,l**) and were correspondingly highly correlated on both cell (**Extended Data Fig. 8c**) and gene (**Extended Data Fig. 8d**) loadings, indicating that they correspond to the same program. Moreover, other factors that differed between Ultima and Illumina were highly related—for example, factor 5 in the Ultima dataset was roughly decomposed into factors 5 and 11 in the Illumina dataset. Overall, we conclude that there is a high correspondence between cell states in Ultima and Illumina data.

As a final test, we evaluated performance with a Perturb-seq screen, where heavy sequencing requirements are particularly limiting for scale ^5, 6^, and used a design that also tested for CITE-Seq ^20^ and Cell Hashing ^21^ performance. Specifically, we used a library from a pilot screen of an ongoing genome-wide Perturb-seq study (PIT, KGS, CJF and AR, unpublished results) to identify regulators of MHC Class I in melanoma A375 cells (**Fig. 4a**). In this pilot, we introduced 6,127 guides targeting 1,902 transcription factors and chromatin modifiers (**Supplementary Table 4**) along with both intergenic and non-targeting control guides, enriched for cells with low HLA levels, and followed by scRNA-seq of 20,000 cells that included CITE-seq ^20^ and Cell Hashing ^21, 22^ (**Online Methods**). We sequenced the resulting scRNA-seq libraries with Illumina and Ultima, but the dial-out libraries used for guide detection, CITE-seq, and Cell Hashing were only sequenced with Illumina (**Extended Data Fig. 9a**-**c**). Initial pre-processing of the Perturb-seq scRNA-seq data showed similar performance for Ultima and Illumina, after sampling reads to have the same number of UMIs in each dataset (**Extended Data Fig. 5a,b, Supplementary Table 1**), as before, as well as in terms of cell assignment to guides (**Fig. 4b** and **Extended Data Fig. 9c**), Cell Hashing barcodes (**Fig. 4c**), and cell clustering and marker gene expression (**Extended Data Fig. 9d**-**i**).

**Fig. 4.**
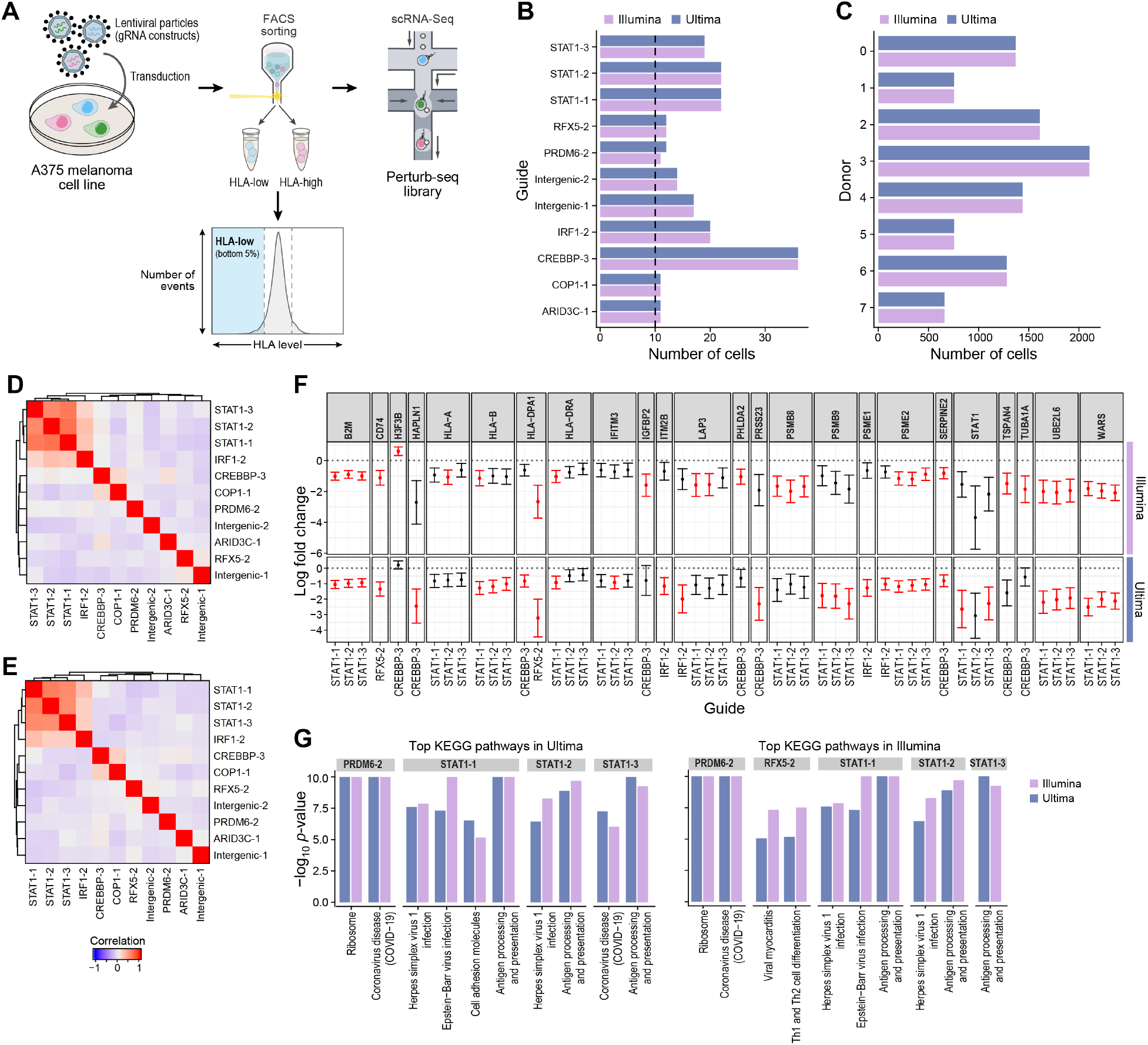
Perturb-seq. (**a**) Perturb-seq to find regulators of MHC Class I in melanoma. We transduced A375 melanoma cells with a genome-wide library, and cells with low HLA expression were enriched by flow cytometry prior to scRNA-seq. (**b**) Number of cells with each perturbation, only plotting those with >10 cells, excluding non-targeting and background guides. (**c**) Number of cells with each Cell Hashing label. Guide similarity heatmaps in Ultima (**d**) and Illumina (**e**). Effects of each guide on each gene in the Illumina data were calculated with an elastic net-based approach as in MIMOSCA. The matrix of guides by genes with (uncorrected) p-values < 0.05 was extracted; the correlation between guides was calculated and plotted as a heatmap. (**f**) We extracted all gene/guide pairs from our DE analysis with FDR < 0.05 in either Illumina or Ultima. For each of these guide/gene pairs, we plotted the logFC on the y axis (with 95% CI) and guide on the x axis, with each box being a different gene, for both Illumina and Ultima. We included all guides that targeted a gene with any significantly different guides. The dots are colored red if significant (FDR < 0.05) and black if not. (**g**) KEGG enrichment analysis for each guide. The 10 pathways with smallest p-values in both Illumina (left) and Ultima (right) are plotted as their -log_10_ p-values in both Illumina and Ultima. All pathways were significant in both Illumina and Ultima at FDR of 0.05.

Importantly, the Ultima and Illumina datasets identified similar relationships between perturbations and similar regulatory effects. For this analysis, we included the 335 cells in Illumina and 336 cells in Ultima, coming from 11 perturbations and 10 control guides in this pilot screen that were assigned to a single perturbation that had more than 10 assigned cells (the same perturbations were found by Illumina and Ultima). We then fit a regularized linear model (with elastic net, similar to ^6, 22^ (**Online Methods**)) of the mean impact of each perturbation on each gene, selected genes with nominal p-values < 0.05 using a permutation-based approach (**Online Methods**), and clustered the guides by these regulatory profiles. Ultima (**Fig. 4d**) and Illumina (**Fig. 4e**) based analyses yielded very similar guide relationships, both between multiple guides to the same gene (*e.g*., *STAT1* guides) and between guides to different, functionally-related genes (*e.g*., *STAT1* and *IRF1* or COP1_1 and CREBBP_3). Moreover, there was very high agreement in the effects on individual genes in both datasets, when comparing DE genes between each guide and an intergenic control (intergenic_1) in each dataset, in both significance and effect size (**Fig. 4f, Extended Data Fig. 9j,k**). Many such gene/guide pairs were significant in both the Ultima and Illumina datasets (**Fig. 4f**), and those significant only in one, had highly similar effect sizes, showing consistent signal. Moreover, the KEGG pathways enriched with DE genes between each guide and an intergenic control were highly similar between the datasets (**Fig. 4g**).

In conclusion, the two sequencing platforms generally perform similarly for scRNA-seq, across two main protocols for droplet based scRNA-seq (3’ and 5’), two different sample types (primary cells and a cell line), and multiple experimental designs (simplex and multiplex, Perturb-Seq, CITE-Seq and Cell Hashing). One key explanation for the minor differences we observed is the position of reads relative to annotated gene boundaries (**Extended Data Fig. 3d**-**g**), as a consequence of Ultima’s single-end reads being closer to gene ends. Additionally, we currently recommend 5’ over 3’ libraries, given the small penalty in lost reads in 3’ libraries (**Fig. 2a**) due to lower sequencing quality adjacent to the poly(T) sequence (**Extended Data Fig. 1**).

It is exciting to imagine what will be possible with Ultima’s lower cost, estimated at over 5-fold reduction in cost per read, that enables sequencing more reads, cells, and/or samples in the context of large scale tissue atlasing projects, such as the Human Cell Atlas ^1^, the BRAIN Initiative ^23^, the Cancer Moonshot Human Tumor Atlas Network ^2^ as well as perturbation screens ^5, 6^. It should also be possible to design droplet-based scRNA-seq reagents, and methods for other large scale single cell and spatial genomics ^24-28^ customized to Ultima sequencing to directly generate libraries and eliminate the need for library conversion (**Fig. 1b**)Additionally, with longer Ultima reads or different construct designs, sequencing of other library types including TCR/BCR and dial-out can be enabled. Finally, such reduced sequencing costs could open the way to use scRNA-seq in clinical applications, including diagnostics (as in next-generation blood tests, “CBC2.0” ^1^) or for therapeutics screens of small molecules, antibodies or cell therapies, impacting both basic biological discoveries and their clinical translation.

## Acknowledgements

We thank Elena Torlai Triglia and Orr Ashenberg for advice on TCR/BCR analysis, Devan Phillips for BL2^+^ room training, and Anna Hupalowska and Leslie Gaffney for help with illustrations and figures. This work was supported by the Klarman Cell Observatory, Broad Institute of MIT and Harvard (A.R., J.Z.L.), Howard Hughes Medical Institute (A.R.), Stanley Center for Psychiatric Research, Broad Institute of MIT and Harvard (J.Z.L.), National Human Genome Research Institute (NHGRI) grant (5RM1HG006193-09) (K.G.S.), NIH F32 Ruth L. Kirschstein Postdoctoral Fellowship (5F32AI138458) (P.I.T.), and NHGRI grants R44HG010558 and R44HG011060 (Ultima Genomics).

## Author contributions

Conceptualization: J.Z.L., A.R., G.A., O.R.R.

PBMC experimentation and Illumina sequencing: X.A.

Perturb-seq design, experimentation, and Illumina sequencing: P.I.T., K.G.S., C.J.F., A.R.

Ultima sequencing: F.O., N.I., O.B.

Computational analysis: S.K.S., G.L.Y., with guidance from A.R.

Supervision: J.Z.L., D.L., A.R.

Writing – original draft: S.K.S., J.Z.L., D.L., G.L.Y., P.I.T., K.G.S., A.R.

Writing – review & editing: everyone

## Data and code availability

RNA-seq data generated in this project will be available on June 1, 2022 from the Gene Expression Omnibus with accession number GSE197452 and the Single Cell Portal at https://singlecell.broadinstitute.org/single_cell/study/SCP1759. The code used for processing and analyzing our sequence data will be freely available on June 1, 2022 from https://github.com/seanken/CompareSequence.

## Methods

### PBMC library preparation

All biospecimens were collected with informed consent by a commercial vendor. Use of all deidentified biospecimens for sequencing at the Broad Institute was further approved by the Broad’s Office of Research Subject Protection (ORSP), which determined that the research did not involve human subjects according to U.S. federal regulations (45CFR46.102f) – determination ORSP-3635. This study complied with all relevant ethical regulations.

We purchased 9 cryopreserved human PMBC samples (AllCells). We thawed PBMC vials in a 37°C water bath for ∼2 minutes. A quick counting revealed high viability (>90%) in all samples. We added 1 ml of RPMI1640 (Thermo Fisher Scientific, 11875093) with 10% Fetal Bovine Serum (Thermo Fisher Scientific, 16140-071), transferred the cells to 15 ml conical tubes, and then added another 9 ml of this media slowly dropwise. We spun down the samples for 10 minutes at 300g at room temperature. After supernatant removal, we flicked each tube to dislodge the pellet and carefully added 10 ml of media dropwise, followed by another spin under the same conditions. After supernatant removal, we flicked the tubes to dislodge pellets, re-suspended the cells in 500 µl PBS 0.4% BSA (Sigma, B8667), and transferred to 1.5 ml tubes. We then spun down the samples for 5 minutes at 300g at room temperature. We washed the cells with 500 µl PBS 0.4% BSA an additional 2 times and filtered through 40 µm cell strainers (Falcon, 352340). We counted cells with a TC-20 cell counter (Bio-Rad) and observed high viability (>90%) for all samples.

For one sample with matched 5’ and 3’ libraries, we loaded one channel of 10x 3’ V3.1 (10x Genomics, 1000128) onto a G chip (10x Genomics, 1000127) and one channel of 5’ V2 (10x Genomics, 1000265) onto a K chip (10x Genomics, 1000286), respectively, aiming to recover 7,000 cells from each. With the other 8 samples, we pooled them equally and loaded onto one channel of 10x 5’ V2 assay aiming to recover a total of 24,000 cells. We generated the 10x 3’ and 5’ scRNA-seq libraries following the manufacturer’s protocols, as well as the TCR and BCR libraries from the 5’ assay with the 10x Chromium Single Cell Human TCR (10x Genomics, 1000252) and BCR Amplification kits (10x Genomics, 1000253), respectively. We performed each experiment once (n = 1 biological replicate). The two 5’ experiments are biological replicates for each other in some, though not all, ways.

### Perturb-seq screen

To generate a large Perturb-seq library targeting all transcription factors and chromatin regulators, we designed a 5706 guide library targeting 1902 genes identified as either transcription factors or chromatin regulators with three gRNAs per gene taking sequences from the Broad Institute Genetic Perturbation Platform Web sgRNA Designer (https://portals.broadinstitute.org/gpp/public/analysis-tools/sgrna-design) ^29^. We included two different types of control gRNAs either guides that cut in a non-gene region (intergenic control) or guides that do not bind any genomic region (non-targeting control) each at 5% of the total guide count. The pooled CRISPR library was cloned as previously described in the CROPseq mKate2 vector backbone ^22^. We transduced Cas9-expressing A375 cells (ATCC CRL-1619) with a transcription factor and chromatin regulator-gRNA library (**Supplementary Table 4**). We selected perturbed cells for 3 days using 2 µg/ml puromycin. After selection, we treated cells with 2 ng/ml recombinant IFNγ for 16 hours. Following IFNγ treatment, we stained cells with CITE-seq and hashing antibodies as previously described ^22^ (**Supplementary Table 4**) along with a fluorescent HLA antibody (BioLegend 311415). The 5% lowest expressing HLA cells were selected via FACS and 40,000 cells were loaded onto one 10x 3’ V3 Chromium channel. We performed this experiment once (n = 1 biological replicate).

### 10x Chromium Illumina sequencing

We sequenced the PBMC libraries on Illumina NextSeq 500 flowcells with at least 20,000 reads/cell for scRNA-seq libraries and 5,000 reads/cell for TCR and BCR libraries. For 3’ libraries, we sequenced 28 bases for Read 1, 55 bases for Read 2 and 8 bases for Index 1. For 5’ libraries, we sequenced 26 bases for Read 1, 45 bases for Read 2 and 10 bases each for Index 1 and Index 2. For TCR and BCR libraries, we sequenced 26 bases for Read 1, 90 bases for Read 2 and 10 bases each for Index 1 and Index 2. We sequenced the Perturb-seq library on Illumina HiSeq X flowcells with 14,000 reads/cell for scRNA-seq libraries, 5,000 reads/cell for CITE-seq libraries, and 1,000 reads/cell for Hashing libraries. For Perturb-seq libraries, we sequenced 28 bases for Read 1, 96 bases for Read 2 and 8 bases for Index 1.

### 10x Chromium library conversion and Ultima Genomics (UG) sequencing

Our 10x Chromium libraries were converted using a library conversion PCR workflow (**Fig. 1b**) to enable sequencing on the UG platform. In brief, library concentration was verified using Qubit (Thermo Fisher Scientific), with conversion PCR library input being 7 ng. Conversion was facilitated through two overhang primers. Primer 1 anneals in the Read 1 region of the 10x library and contains a UG specific overhang (Index Adapter sequence – IA). It contains primer binding sites for clonal amplification and sequencing. IA also includes an in-line UG-specific sample barcode (PS-SBC). Primer 2 anneals in the Read 2 region and contains a UG-specific overhang (Unique Bead Adapter sequence – UBA) necessary for clonal amplification (**Supplementary Table 5**). We used the Q5 Hot Start High-Fidelity kit (New England Biolabs) with 10 PCR cycles for amplification, followed by DNA Clean Concentrator tubes (Zymo Research) as per the manufacturer’s instructions for PCR product purification, and quantification of the purified library.

After pooling, we seeded libraries, clonally amplified them on sequencing beads using a high-scale emulsion amplification tool, and sequenced them on a prototype UG Sequencer, which uses a sequence of additions of partially labeled non-terminating nucleotides, followed by imaging to generate single-end reads of length 200 to 250 bases (**Fig. 1c**, a detailed workflow description will be published elsewhere). For sequencing of 10x Chromium 3’ libraries, we used a modified sequencing protocol that accommodates the high consumption of dT nucleotides in the poly(dT) stretch of the cDNA. Specifically, we included additional T injections when sequencing cycles 28 to 32, which were predicted to include the poly(dT) stretch: (TGCA)_27_ (T_10_GCA)_5_ (TGCA)_60_.

### Initial Ultima read processing

To enable standard scRNA-seq analysis of single-end reads, we first converted UG data to create paired-end data (**Fig. 1d**). To this end, we removed conversion adapters, and quality trimmed reads using Cutadapt v2.10 ^30^, using a threshold of 30. We discarded reads not containing at least 8 Ts for the expected poly(T) for 3’ libraries or the template switch oligo (TSO) for 5’ libraries. We split reads into two sequences: one containing the CBC and UMI, and the other containing the reverse complement of the cDNA using Cutadapt and SeqKit v0.15.0 ^31^. To create Read 1 (<output_read1>) and Read 2 (<output_read2_revcom>) files from input FASTQ files, we ran the following three commands. For 3’: cutadapt -j 0 --discard-untrimmed --pair-filter any -a CTACACGACGCTCTTCCGATCT;max_error_rate=0.2;min_overlap=10;required…AGATCG GAAGAGCACACGTCTG;max_error_rate=0.2;min_overlap=6 -U 50 -q 30 -A TTTTTTTTTTTT;max_error_rate=0.2;min_overlap=8;required…AGATCGGAAGAGCACAC GTCTG;max_error_rate=0.2;min_overlap=6 -o <output_read1_long> -p <output_read2> -- minimum-length 28:50 <input_fastq> <input_fastq>

cutadapt -j 0 --minimum-length 28 --maximum-length 28 --length 28 -o <output_read1> <output_read1_long>

zcat <output_read2> | awk ‘{if ((NR%4 == 2) || (NR%4 == 0)) {print substr($0,1+5,90) } else

{print $0 } }’ | seqkit -j 8 seq -p -r -t DNA | gzip > <output_read2_revcom>

For 5’: cutadapt -j 0 --discard-untrimmed --pair-filter any -a CTACACGACGCTCTTCCGATCT;max_error_rate=0.2;min_overlap=10;required…AGATCG GAAGAGCACACGTCTG;max_error_rate=0.2;min_overlap=6 -U 48 -q 30 -A ^TTTCTTATATGGG;max_error_rate=0.5;min_overlap=8;required…AGATCGGAAGAGCACACGTCTG;max_error_rate=0.2;min_overlap=6 -o <output_read1_long> -p <output_read2> -- minimum-length 26:50 --maximum-length 390:315 <input_fastq> <input_fastq>

cutadapt -j 0 --minimum-length 26 --maximum-length 26 --length 26 -o <output_read1>

<output_read1_long>

zcat <output_read2> | awk ‘{if ((NR%4 == 2) || (NR%4 == 0)) {print substr($0,1+3,90) } else

{print $0 } }’ | seqkit -j 8 seq -p -r -t DNA | gzip > <output_read2_revcom>

We removed reads with a cDNA sequence < 50bp. For 3’ libraries, we clipped the first five bases after the poly(T) and masked the last three bases of the UMI. For 5’ libraries, we clipped the first three bases after the TSO. We trimmed cDNA sequences to 90 bases.

### Extracting expression information from FASTQ files

We used Cell Ranger v5 ^14^ to pre-process data for both Illumina and Ultima (for Ultima using simulated Read 1 and Read 2 as extracted above). For all datasets, we used the GRCh38 human reference from 10x Genomics unless otherwise stated, and set --expect-cells to the expected number of cells (7,000 cells for the single sample 3’ and 5’ PBMC data, 24,000 cells for the 5’ mixture PBMC sample, and 20,000 cells for the Perturb-seq sample). To process 3’ Ultima data, unless otherwise stated, we modified the last 3 bases of the UMI using awk by replacing them with A’s and setting the last 3 quality values to be equal to I.

For the 5’ mixture data, we demultiplexed it by first calculating SNP coverage data for SNPs in the 1000 Genomes Project ^32^ with cellsnp-lite v1.2.0 ^33^ (using --minMAF 0.1 --minCOUNT 20) followed by Vireo v0.5.5 ^16^ to get labels for the sample of origin.

For Perturb-seq data, we also passed Cell Ranger FASTQ files for dial-out data (using the CRISPR Guide Capture keyword), hash tag oligo (HTO) data (using the Custom keyword), and antibody derived tag (ADT) reads for CITE-seq (using the Antibody Capture keyword), as well as feature barcode information for each. We further processed HTO data with DemuxEM v0.1.7 ^34^ to obtain sample labels.

### Sampling reads

For each Ultima dataset, unless otherwise stated, we sampled it to have both the same number of reads and the same number of total UMI as the corresponding Illumina dataset. This was performed by sampling the FASTQ files with seqtk v1.0 sample (https://github.com/lh3/seqtk) passing it the argument -s 100, the FASTQ file to sample, and the proportion to sample by. For sampling to the same number of reads, we calculated the proportion by dividing the total number of reads in Illumina by the number in Ultima. For sampling to the same number of UMIs, we used DropletUtils v1.10.3 ^35^ to sample Ultima data to different levels (in 5% increments) and calculated the total number of UMIs. We chose an initial unrefined proportion as the largest proportion that gave fewer UMIs in Ultima than were present in Illumina. We then performed a refinement step with 1% steps ranging from this unrefined proportion up to the initial unrefined proportion plus 5%, and chose the final refined proportion used for sampling from this range to be the largest proportion that gave fewer UMIs in Ultima than were present in Illumina. We did not sample the 5’ mixture Ultima data because it had roughly the same number of UMIs as the 5’ mixture Illumina data. After sampling, we processed data in a similar fashion to non-downsampled data (see **Extracting expression information from FASTQ files**).

### Extracting FASTQ QC metrics

To extract base quality information from each FASTQ file, we randomly selected 1,000,000 reads with seqtk sample using the parameter -s 100 to set a random seed. We then used the SeqIO.parse function from Biopython v1.79 ^36^ to read the FASTQ into Python. We then extracted the quality information with the letter_annotations function and recorded the resulting information to a file with one line per read and one column for each base in that read. This was used for downstream visualizations.

To explore the effects of shortening UMIs on number of reads, we loaded the molecular information h5 file generated by Cell Ranger into R with DropletUtils and saved the resulting data frame. We then loaded this into Python, resulting in a table with one entry for each UMI counted by Cell Ranger, which included CBC, UMI, and Gene. We shortened the UMI to different lengths and used the pandas ^37^ groupby function to count the number of UMIs that collapsed together after this shortening and the number of UMIs from different genes that collapsed together after this shortening.

### Analysis of PBMC data

We loaded filtered PBMC count data from Cell Ranger (located in the outs/filtered_feature_bc_matrix subdirectory output) into R v4.0.3 ^38^ using Seurat v4.0.0 ^39^. To avoid biases introduced by using slightly different sets of cells, we used only the intersection of the sets of cells found in Ultima and Illumina for each analysis. For the 5’ mixture data, we also uploaded the labels from Vireo and removed doublets and unassigned cells. We processed data were then processed through the standard Seurat pipeline, as followed. We scaled scaling data to transcripts per million (TPM) and log normalized with NormalizeData, finding variable genes with FindVariableFeatures (using 2,000 variable genes), scaling data and regressing out the number of genes per cell with ScaleData, and performing PCA with RunPCA. We performed UMAP embedding and clustering using Seurat’s FindNeighbors followed by FindClusters with 20 Principal Components (PCs) and otherwise default parameters (including FindClusters using Louvain clustering with a resolution of 0.8). Cell types were assigned using Azimuth v0.3.2 ^39^ with the built-in PBMC dataset. In particular, we assigned cell types at two different levels of granularity, denoted in the Azimuth labeling by l1 (general cell types) and l2 (refined cell types). Adjusted Mutual Information (AMI) was calculated with the AMI function in the aricode package v1.0.0 (https://github.com/jchiquet/aricode). For calculating joint embeddings, we combined and processed the two datasets through the same Seurat pipeline, except the sequencing technology used (Ultima or Illumina) was regressed out before PCA with ScaleData. Presto v1.0.0 ^17^ was used to calculate DE genes between cell types with default parameters using an FDR cutoff of 0.05.

### Analysis of method specific biases

We calculated logFCs comparing expression levels between Ultima and Illumina by creating a pseudobulk profile for an entire dataset using count data, which was then normalized to TPM. We calculated logFC values by taking the difference between the log of the corresponding TPM values in Ultima and Illumina with a pseudocount of 10. All plots of log TPM have a pseudocount of 1 added.

For the sampling analysis to compare reads with each other from the same sequencing run, we used a modified version of the downsampleReads code from DropletUtils applied to the molecule_info.h5 file from Cell Ranger to split reads into two disjoint sets of equal size (up to rounding) and calculated the associated gene by cell UMI count matrix for each set of reads.

We performed DE analysis between Ultima and Illumina with Presto ^17^. To explore GC and length biases in DE results, we used a Python script to process the GTF file used in Cell Ranger and collapse overlapping exons from the same gene into one genomic interval. This information was written to a BED file and used to calculate the total length of sequence in each gene covered by at least one exon (the gene length used in our analysis). We used the “bedtools getfasta” command in BEDTools version 2.26.0 ^40^ with arguments -s -name -tab to extract the associated sequences of each of the regions in the above BED file. We processed the resulting FASTA file with a Python script that extracted gene level GC information.

Classification of gene expression and read assignment bias was performed as follows. For each protein-coding gene with a total count >100 in at least one of the platforms, we calculated read assignment rate (see below) differences for unfiltered (total reads, including reads from cell barcodes not in the filtered list) Illumina and downsampled Ultima reads, and normalized the fold-changes of counts by the median difference. We partitioned the genes into three categories: (1) >5-fold higher in Illumina (“Higher count in Illumina”); (2) >5-fold higher in Ultima (“Higher count in Ultima”); and (3) all remaining genes (“Similar counts”) (**Extended Data Fig. 3c**). A read assignment rate higher in Illumina (“Higher read assignment in Illumina”) was defined if the ratio of Cell Ranger gene-assigned reads out of the total reads mapped to the annotated gene body plus a flanking 100bp was higher in Illumina (>3-fold higher ratio with p < 0.01, binomial test). The read assignment rate higher in Ultima (“Higher read assignment in Ultima”) was defined analogously.

To explore 3’ and 5’ bias, the GTF file used by Cell Ranger was processed as described above to collapse overlapping exons together. We then used pysam v0.15.3 (https://github.com/pysam-developers/pysam) to load each alignment (selecting 1% of alignments at random), excluding those without an assigned gene, CBC, or UMI, as well as excluding multimappers. We then calculated the distance along the exonic regions of the gene (normalized by gene length) from the 5’ end of the gene to the 3’ and 5’ ends of the read using the overlapping exon representation we generated earlier. We then recorded this information for each alignment. For plotting, this was divided into bins of length 1% labeled from 0 to 100 (including the bottom of each bin but not the top – meaning that bin 100 was empty for 5’ reads and bin 0 was empty for 3’ reads), and normalized by the number of reads mapping to that gene to avoid highly expressed genes biasing the results.

To extract information about the number of reads falling into different categories (unmapped, ambiguous, etc.), we took the BAM file from Cell Ranger and applied FeatureCount v1.6.2 ^41^ with settings -t exon -g gene_name –fracOverlap 0. For 3’ data we set -s 1 to denote sense reads, while for 5’ data we used -s 2 to denote antisense reads. In addition, for gene level information about the number of reads in different categories, we reran FeatureCount once with the flag -M (for multimappers) and once with the flag -O (for ambiguous reads).

For IGV v2.3.80 ^42^ plots, we used SAMtools v1.8 ^43^ to extract a region around each gene of interest from the associated 10x BAM file. We then used grep to extract reads that were assigned to a given gene and those that were not.

### Extension of the standard reference

We built a Nextflow-based ^44^ pipeline that takes in a preexisting reference GTF file and RNA-seq BAM file (from paired-end or single-end RNA-seq) and outputs a new GTF file that extends the old one using RNA-seq data. The first step in the pipeline annotates reads in the BAM file overlapping genes in the standard reference using FeatureCount with the parameters -t exon, -R BAM, and -g gene_id, as well as using the -s 1 or –s 2 flag depending on the strandedness of the RNA-seq data. For paired-end data we also used the flag -p. We then used a pysam-based start and end coordinates being the start and end coordinates of that read, and with an extra field recording the assigned gene for the read. We excluded reads with large gaps (>10 bases labeled as N in the CIGAR string) and, for paired-end reads, only include properly paired reads. We then sorted the BED file with bedops ^45^, clustered the entries in this BED file using bedtools cluster with the -s flag, and used bedtools groupby to merge BED entries from the same cluster and gene. We then sorted the resulting BED file with bedops again, use a Python script to turn the BED file into a GTF with one exon per entry in the BED file. We combined this GTF file with the GTF for the standard reference and sorted the results with BEDTools, yielding the extended references. These new GTFs were then used to generate references for Cell Ranger with cellranger mkfastq. We used this approach to create three references, one using published bulk data ^12^ (using the BAM file generated in that publication), and two using scRNA-seq Ultima PBMC data – one generated with 3’ data and the other with 5’ data. We then processed PBMC data with each of these references using cellranger count as described above and performed downstream analysis. We did not process the 3’ data with the 5’ reference or vice versa.

### Processing TCR and BCR data

We processed FASTQ files for 10x Chromium TCR and BCR data with Cell Ranger v5 using the vdj command and the prebuilt 10x reference (refdata-cellranger-vdj-GRCh38-alts-ensembl-5.0.0). We then loaded data into the associated Seurat object with the djvdj package v0.0.0.9000 (https://rdrr.io/github/rnabioco/djvdj/man/djvdj-package.html) where they were used for downstream visualizations.

### Gene program analysis by NMF

We calculated all NMF models with RcppML v0.3.7 ^46^ using 15 factors and with the log TPM matrix as input including genes expressed in more than 1% of cells. NMF returns a cell loading matrix, with one row per cell and one column per factor, and a gene loading matrix, with one row per gene and one column per factor. To test how well NMF factors from one data type fit another, we split our data into a training set (with 5,000 cells) and a test set (all other cells) to avoid data overfitting when testing the gene loading matrix. We then fit NMF models separately on the Ultima data, Illumina data, and a permuted version of the Illumina data (where the values of each gene were randomly scrambled between cells) using the 5,000 cell training set. To test the accuracy of gene loadings of each NMF model for each data type, we used the project function from RcppML to generate a cell loading matrix on the training data and the mean squared error (MSE) was measured. For cell loadings, we used a similar approach, except that testing was performed on the test dataset. In all cases we repeated the analysis 10 times with different random seeds to account for variability in NMF solutions.

For consensus NMF (cNMF) ^19^, a modification of NMF meant to improve robustness and reproducibility, we implemented an R version with two minor changes: (1) Instead of performing NMF on the count data with each gene normalized by its standard deviation, we performed it on log TPM data; and (2) we performed each NMF round used by cNMF with the variable genes from our Seurat analysis instead of the variable gene selection procedure in the cNMF package. We ran cNMF on all cells in each dataset, using 100 iterations of NMF and 15 factors. For Perturb-seq based cNMF, we used the project function in RcppML to project the gene loadings onto the PBMC mixture dataset. Similarity matrices were calculated with Pearson correlation.

### Analysis of Perturb-seq data

We processed Perturb-seq data through a similar pipeline to the PBMC data (see **Analysis of PBMC data**), except the HTO, ADT, and guide count matrices were also uploaded as additional assays, while the DemuxEM labels for Hash ID and the Cell Ranger labels for guide assignment were added to the metadata. After initial processing with Seurat, we removed cells assigned to multiple hash tags or multiple guides, as were cells assigned to guides with 10 or fewer cells assigned to them.

We generated a guide similarity matrix with a slightly modified version of the MIMOSCA package ^6^. We extracted a cell by gene log TPM expression matrix from the Seurat object, selecting the cells filtered as described above (one hashtag and guide assignment, with guides with more than 10 assigned cells) and genes that were expressed in >5% of cells. We also extracted a covariate matrix consisting of the scaled number of UMIs per cell, as well as one-hot encoded versions of the perturbation assignments, Hash ID assignments, and cluster assignments (based on clustering all cells with Seurat’s FindClusters function at a resolution of 0.2 with 20 PCs and otherwise default settings). We loaded data into Python using pandas and an elastic net model was fit modeling expression as a linear model of the covariates using sklearn.linear_model.ElasticNet ^47^ with parameters l1_ratio=0.5, alpha=0.0005, and max_iter=10000. The coefficient matrix from this model was saved. We randomly permuted guide labels 100 times (while preserving the number of guides assigned to each hash barcode and vice versa) followed by the same elastic net-based analysis. We loaded the resulting gene by covariate coefficient matrices into R and discarded columns that did not correspond to guide labels, along with columns corresponding to non-targeting control guides (those labeled as NO_SITE in our feature data) and the Background control guide. For each gene, we calculated a p-value based on the resulting matrices by scoring each gene by the maximum absolute value for that gene across all guides. We partitioned genes into 20 bins of equal size based on average expression. We calculated a p-value for each gene by comparing the score of that gene in the non-permuted data to the score of all genes in the same bin as it in all 100 permuted datasets. We retained all genes in the coefficient matrix with uncorrected p-value < 0.05 and calculated the Pearson correlation between guides based on this matrix.

### Perturb-seq differential expression analysis of genes regulated by each guide

We performed DE between each guide’s profiles and the intergenic guide profiles with Nebula v1.1.7 ^48^, with the assigned Hash ID as the sample of origin, and the Intergenic_1 guide as reference. We calculated Benjamini-Hochberg FDR ^49^ on the resulting p-values. We performed KEGG enrichment analysis with the KEGG ontology ^50^ using ClusterProfiler v3.18.1 ^51^ and performing GSEA with the fgsea package v1.16.0 ^52^. KEGG terms with fewer than 20 genes were filtered before visualization.

### Visualization

Most of the visualization was performed using ggplot2 v3.3.3 ^53^ and cowplot v1.1.1 (https://github.com/wilkelab/cowplot) packages in R. The major exception to this was the heatmaps, which were produced with the ComplexHeatmap package v2.6.2 ^54^ and NMF package ^55^, and histograms that were produced with the base R hist function.

## Extended Data Figures

**Extended Data Figure 1. Sequence read quality for matched 5’ and 3’ PBMC libraries**.

(**a**) Bar plot of the percent of reads at different quality levels for each sequencing platform and library type along the length of the read, for either Read 1, or Read 2 (or, for Ultima, the subsection of the read corresponding to Read 2), or the full read (up to 200 bases, Ultima only). Annotation of the reads is shown at the bottom. Information to guide UMI trimming from the UMI 3’ end for 3’ libraries (**b-d**). (**b**) Rarefaction curves at different UMI lengths and with and without filtering low quality reads. (**c**) At different UMI lengths, UMI/CBC pairs that have all Read 2’s in the same gene. Lower levels suggest UMIs are too short. (**d**) Number of UMIs at different UMI lengths – a measure of similar UMIs collapsing as length decreases.

**Extended Data Figure 2. Quality metrics for matched 5’ and 3’ PBMC libraries sampling reads to the same number of UMIs**.

(**a**) Number of cells identified by Cell Ranger only in Ultima, only in Illumina, or both. (**b**) Distribution of the number of genes per cell. (**c**) Scatter plots comparing gene expression in Ultima and Illumina sequencing. (**d**) Scatterplots comparing reads sampled from the same sequencing run. For (**c**) and (**d**), one point for each gene as in **Fig 2e**. For all 3’ libraries, the last three UMI bases were trimmed for quality reasons.

**Extended Data Figure 3. Sequencing biases in matched 5’ and 3’ PBMC libraries**.

(**a**) Genes were assigned to bins by GC content (exonic sequences only) and the average logFC for each bin between Ultima and Illumina is shown. Positive logFC indicates higher expression in Ultima. (**b**) Genes were assigned to bins by log_10_ length (total number of bp in at least one exon for a given gene) and the average logFC for each bin between Ultima and Illumina is shown. Positive logFC indicates higher expression in Ultima. (**c**) Differences in read assignment ratio for genes categorized based on their expression being similar or higher in one platform than the other. Number of genes in each category shown on top of each bar. See Materials and Methods for additional details. (**d**) For each library (5’ PBMC, left, and 3’PBMC, right), the 5’ end of each read is shown at relative positions along the length of each gene. (**e**) Relative position of the 3’ end of reads as in **(d)**. (**f**) *LILRA5* sequence coverage of 3’ libraries with Illumina (top two tracks) and Ultima (next two tracks). Standard gene annotation with introns as lines, exons as boxes, and arrows for direction of transcription shown in bottom track. Extended reference annotation using bulk or single cell (SC) Ultima data shown at bottom. Reads assigned (blue) or not (red) to *LILRA5* by Cell Ranger. (**g**) *HIST1H1D* sequence coverage of 5’ libraries shown as in (**f**). (**h**) Bar plots showing classification of reads by FeatureCount. Ultima (short) reads had Read 2 trimmed to 45 bases for 5’ and 55 bases for 3’ to match Illumina read lengths. Assigned: assigned to genes (uniquely mapped reads that overlap a gene; not all of these are assigned to a gene by Cell Ranger, which only assigns reads that completely overlap a gene); ambiguous: uniquely mapped reads that overlap the exons of multiple genes; no feature: uniquely mapped reads that do not overlap any exons; multimappers: map to multiple genomic locations (note Cell Ranger is able to recover multimappers if they only overlap one gene); unmapped: map to no genomic locations. Each read can only fit in one category. Ultima data sampled to have the same number of UMIs ((**a**) and (**b**)) or reads (other panels) as the Illumina data.

**Extended Data Figure 4. Gene expression and read assignment with extended references**.

(**a**) Scatter plots comparing gene expression of Illumina and Ultima datasets using extended references for both. Extended references from bulk RNA-seq (bulk) or Ultima scRNA-seq (SC). Labeling as in **Fig. 2e**. For 5’ data we used the reference extended by 5’ Ultima data, while for 3’ data we used the reference extended by 3’ Ultima data. (**b**) Same as (**a**), except using the standard reference for Illumina in all cases. (**c**) Same as (**a**), except comparing standard with extended references for the same dataset. (**d**) Bar plots showing classification of reads by FeatureCount, both with the standard reference and with the extended references. Categories as in **Extended Data Fig. 3**. Reads were sampled so that Illumina and Ultima have the same number of reads.

**Extended Data Figure 5. Quality metrics for libraries from a mixture of PBMCs** (**5’) and from Perturb-seq (3’)**.

(**a**) The total number of UMIs detected per cell at different sequencing depths. (**b**) The number of cells found by Cell Ranger only in Ultima, only in Illumina, or in both. (**c**) Distribution of the number of genes per cell. (**d**) Scatter plots with one point for each gene as in **Fig. 2**. For the 3’ library, the last three UMI bases were trimmed for quality reasons. For panels (**b**) to (**d**), reads sampled to have the same number of UMIs for Illumina and Ultima for 3’ libraries, but sampling not needed as already nearly the same for 5’ libraries.

**Extended Data Figure 6. PBMC marker gene and TCR/BCR expression**.

Heatmaps showing expression of known cell type markers ^12^ in Illumina (**a**) and Ultima (**b**) libraries in Azimuth-assigned cell types. We did not use markers for plasma cells or markers selected because they are downregulated in a given cell type. (**c**) Percent of cells in each Azimuth-assigned cell type with associated TCR or BCR clonotype information. (**d**) Top TCR clonotypes among T cells, stratified by individual, colored by more refined cell type in Illumina (top row) and Ultima (bottom row).

**Extended Data Figure 7. Cell type characterization of matched 5’ and 3’ PBMC libraries**. UMAP plots of Azimuth-assigned cell types in 5’ (**a**) or 3’ (**b**) libraries. UMAP plots of Azimuth-assigned cell types on a joint embedding in 5’ (**c**) or 3’ (**d**) libraries. (**e**) Bar plots of cell type proportions for the 5’ libraries. There is 92% agreement in the cell type labels between the two sequencing platforms. (**f**) Bar plots of cell type proportions for the 3’ libraries. There is 96% agreement in the cell type labels between the two sequencing platforms.

**Extended Data Figure 8. Comparisons of cell state for the PBMC mixture between sequencing platforms**.

(**a**) NMF applied to either Illumina, Ultima, or Illumina data with permuted genes, to extract the cell loadings, and test how well they fit either the Illumina (left) or Ultima (right) data, measured by Mean Squared Error (MSE; lower MSE is better). Dots show analysis repeated ten times with different seeds. (**b**) NMF applied to either Illumina, Ultima, or Illumina data with permuted genes, to extract the gene loadings, and test how well those loadings fit either the Illumina (left) or Ultima (right) data, measured by MSE. Dots as in (**a**). (**c**) Correlation between cell level loadings shown after applying cNMF on Illumina and Ultima data with 15 factors. (**d**) Correlation between gene level loadings shown after applying cNMF on Illumina and Ultima data with 15 factors. (**e**) Correlation between cell level loadings of both runs after applying cNMF on the same Illumina data twice with 15 factors. (**f**) Correlation between gene level loadings of both runs after applying cNMF on the same Illumina data twice with 15 factors. (**g**) Correlation between cell level loadings in the Illumina data after applying cNMF. (**h**) Correlation between gene level loadings in the Illumina data after applying cNMF. (**i**) Correlation between cell level loadings after performing cNMF on the PBMC Illumina and Perturb-seq Illumina data with 15 factors and projecting the Perturb-seq gene loadings onto the PBMC data to get cell loadings. (**j**) Correlation of gene level loadings after performing cNMF on PBMC Illumina and Perturb-seq Illumina data with 15 factors. Feature plots of the cell level loading cNMF factors for Ultima (**k**) and Illumina (**l**) in the joint UMAP space. All correlations here are Pearson correlations.

**Extended Data Figure 9. Additional Perturb-seq analysis**.

(**a**) Histogram of the number of guide UMIs per cell from dial-out PCR. (**b**) Histogram of the number of CITE-seq ADT UMIs per cell. (**c**) Histogram showing the number of guides with a given number of cells assigned to them. Note the y axis is log scaled. Dotted line: 10 cells per guide. (**d**) UMAP of gene expression for Illumina. (**e**) Feature plot of the number of genes for Illumina. (**f**) Violin plots of DE genes among clusters in the Illumina data. Shown are clusters showing active cell cycling (cluster 0, 1, and 5) and a cluster with high immediate early gene levels (cluster 5). (**g**) UMAP of gene expression for Ultima. (**h**) Feature plot of the number of genes for Ultima. (**i**) Violin plots of DE genes among clusters in the Ultima data. (**j**) Scatterplot of -log_10_ (p-value) for each guide/gene pair in Illumina vs. Ultima, where the p-values are the output of DE analysis comparing each guide to the control Intergenic_1 guide. (**k**) Scatterplot of logFC for each guide/gene pair as in (**j**). Only gene/guide pairs with uncorrected p-value < 0.01 in either Ultima or Illumina are shown.

## Supplementary Tables

**Supplementary Table 1**. Sequencing metrics.

**Supplementary Table 2**. Comparison of read mapping between Ultima and Illumina.

**Supplementary Table 3**. Differentially expressed genes between Ultima and Illumina with standard or extended references.

**Supplementary Table 4**. Antibody, Perturb-seq, and hashing DNA barcodes.

**Supplementary Table 5**. PCR primers to convert libraries for Ultima sequencing.

