## Extended Data Figures for "Single cell RNA-seq by mostly-natural sequencing by synthesis"

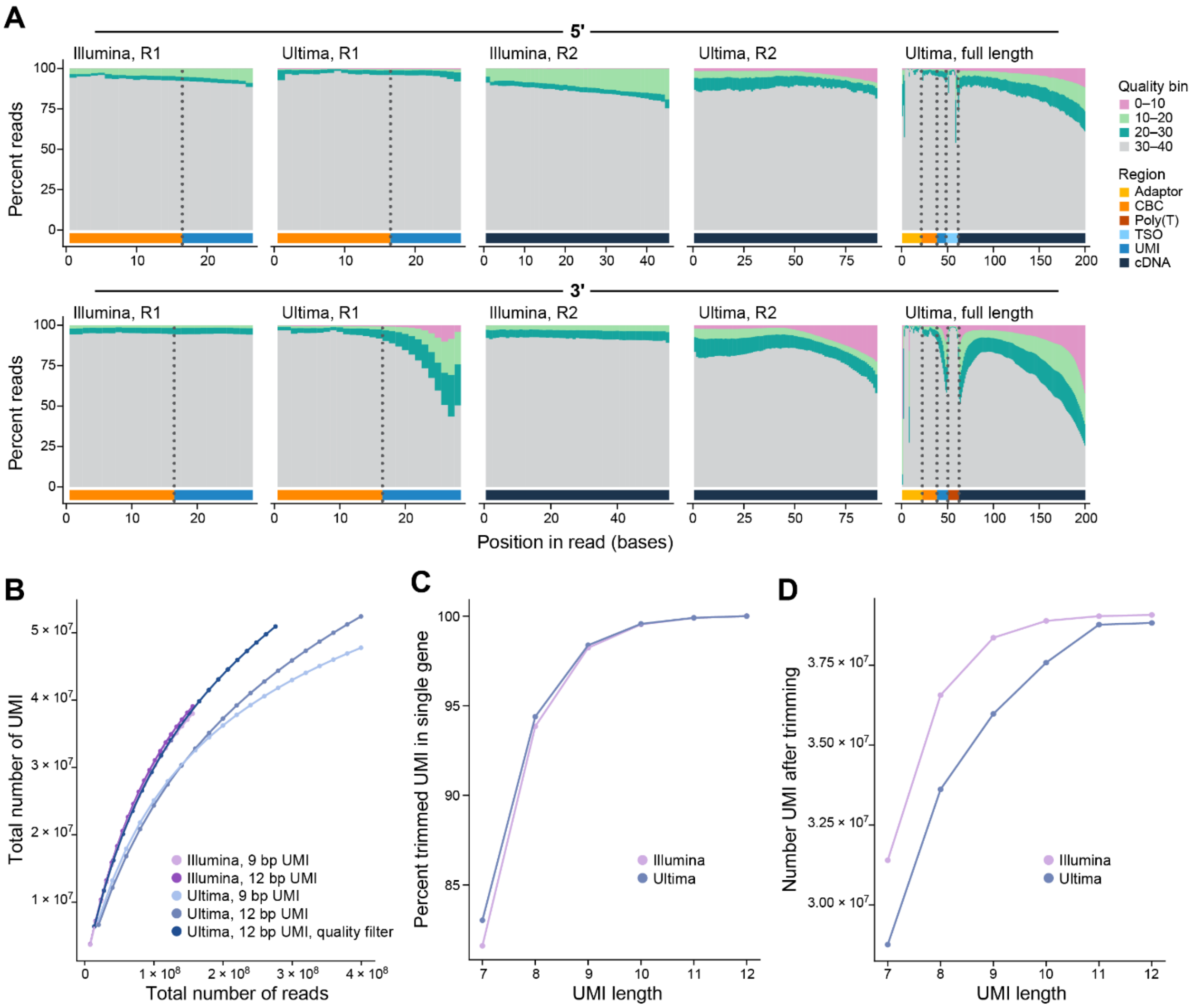


**Extended Data Fig. 1.**

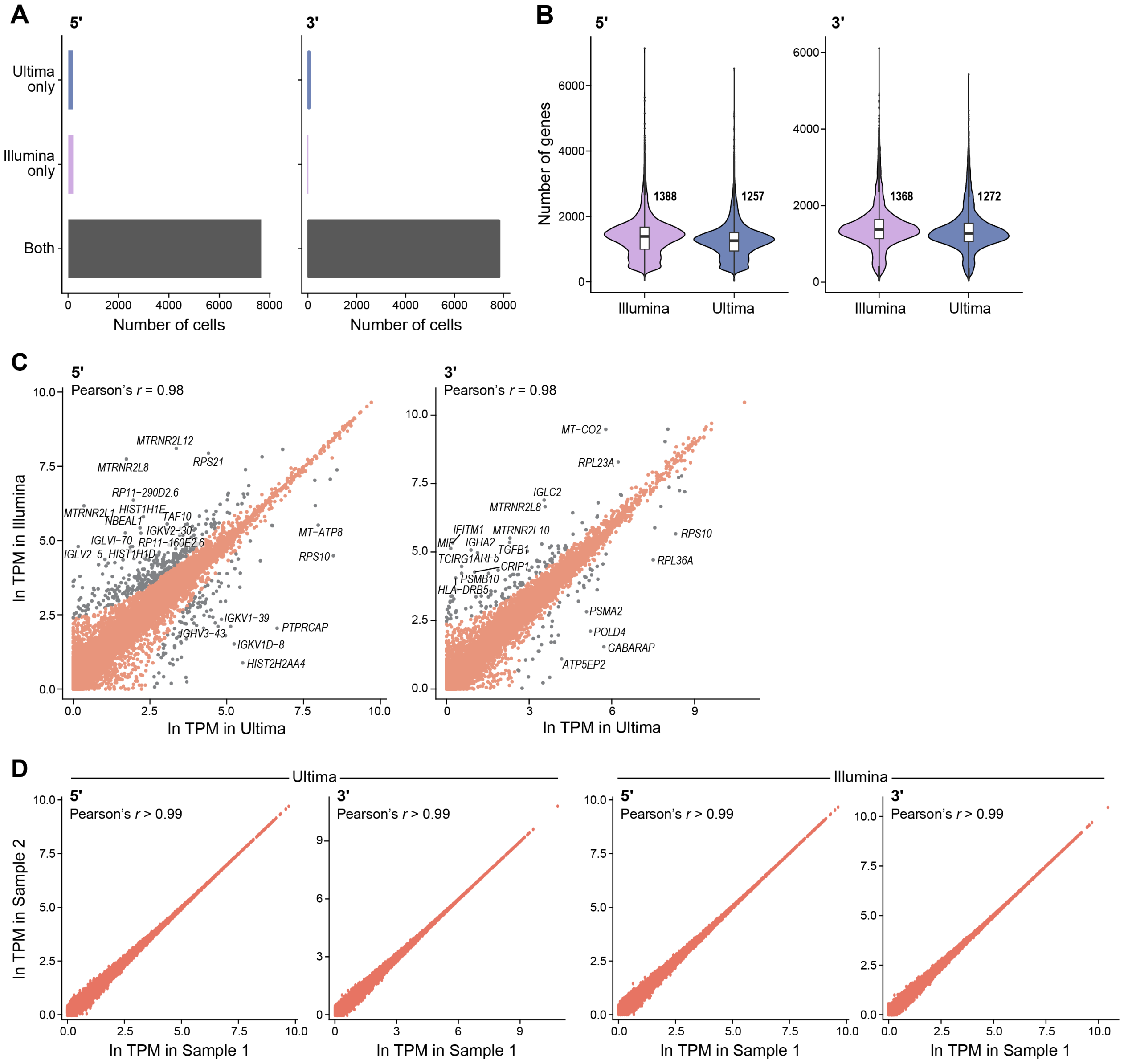


**Extended Data Fig. 2.**

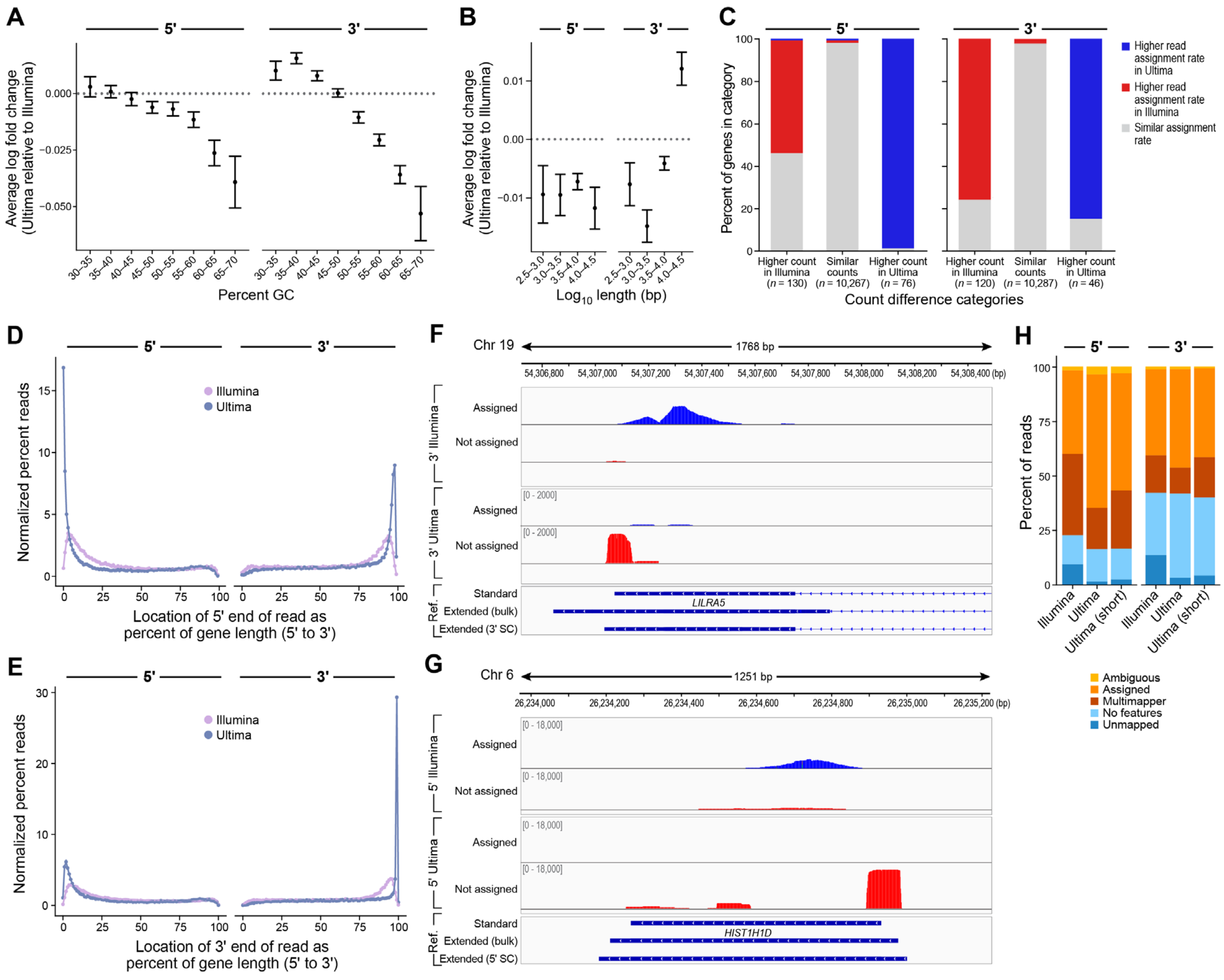


**Extended Data Fig. 3.**

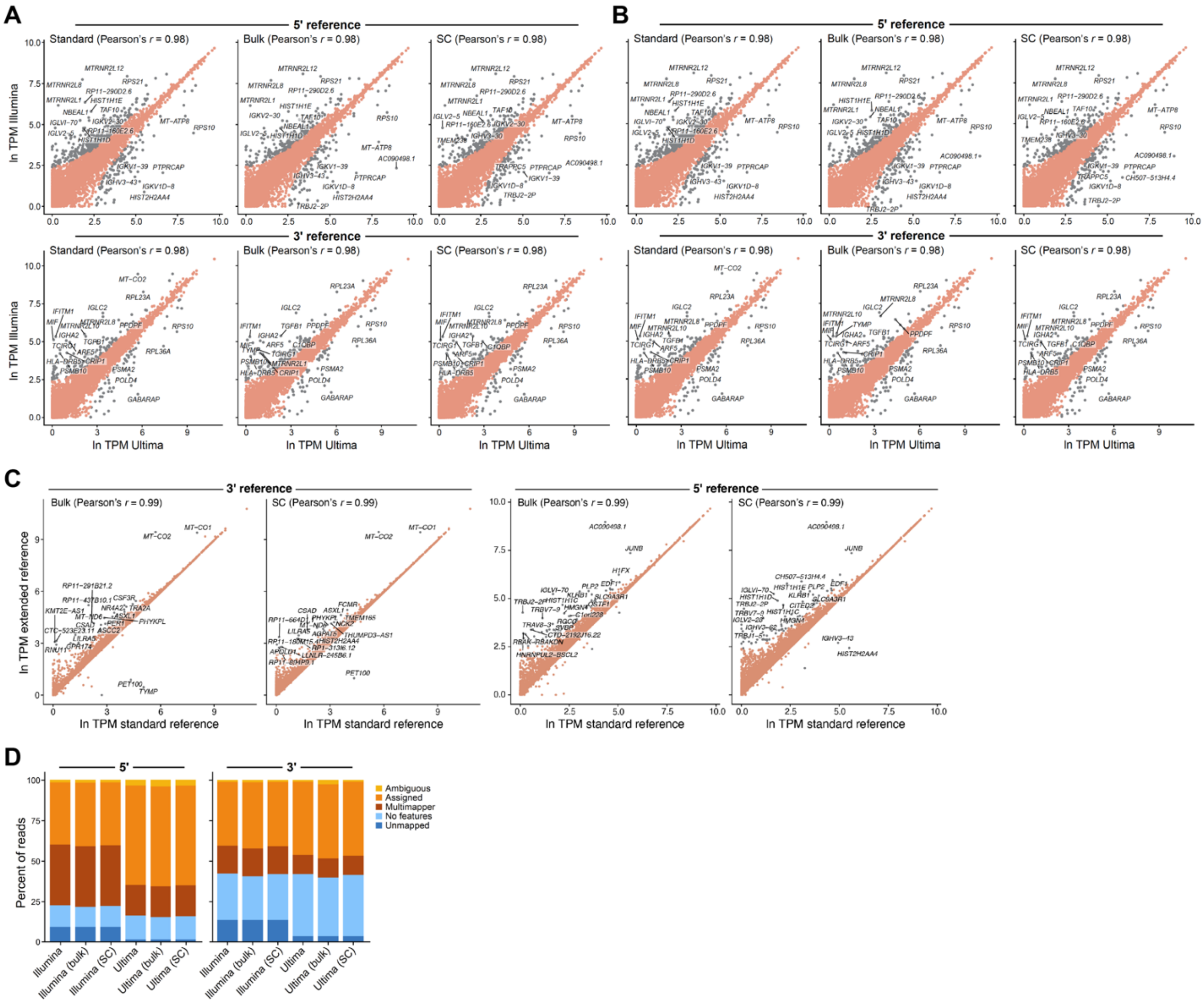


**Extended Data Fig. 4.**

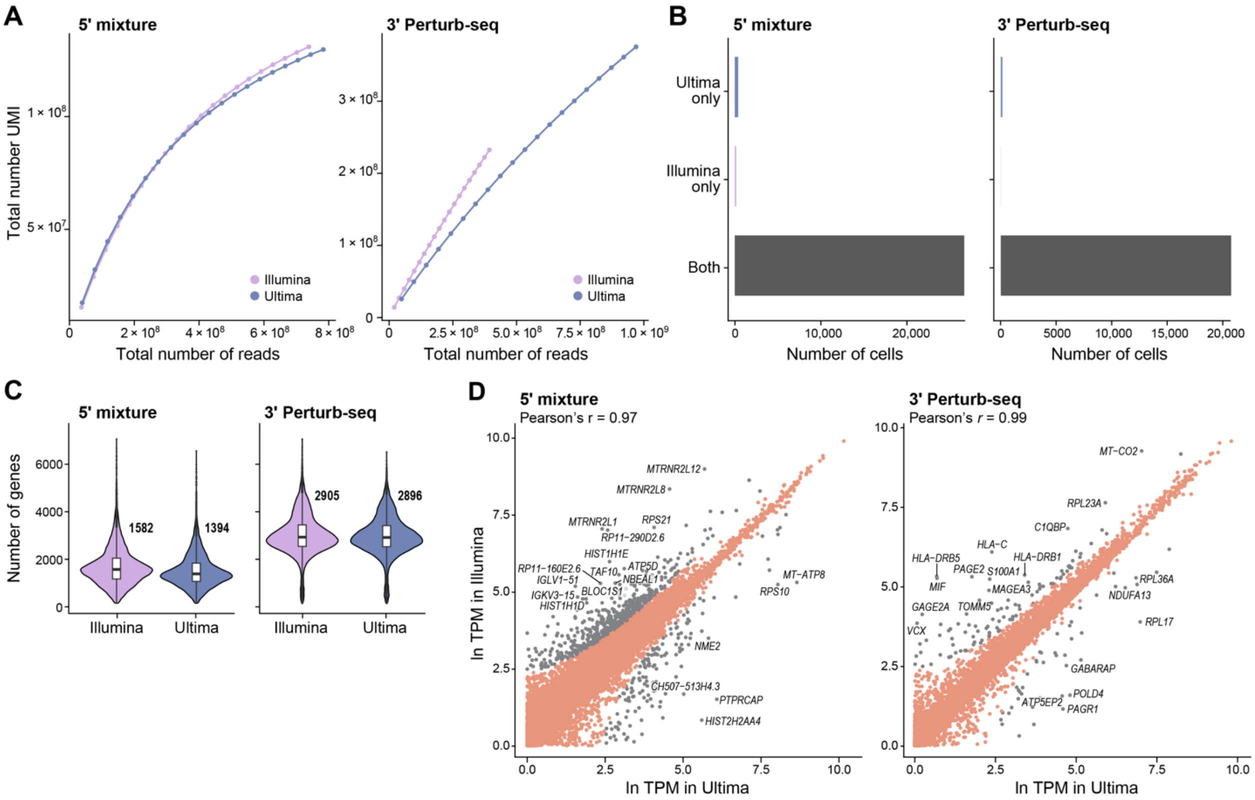


**Extended Data Fig. 5.**

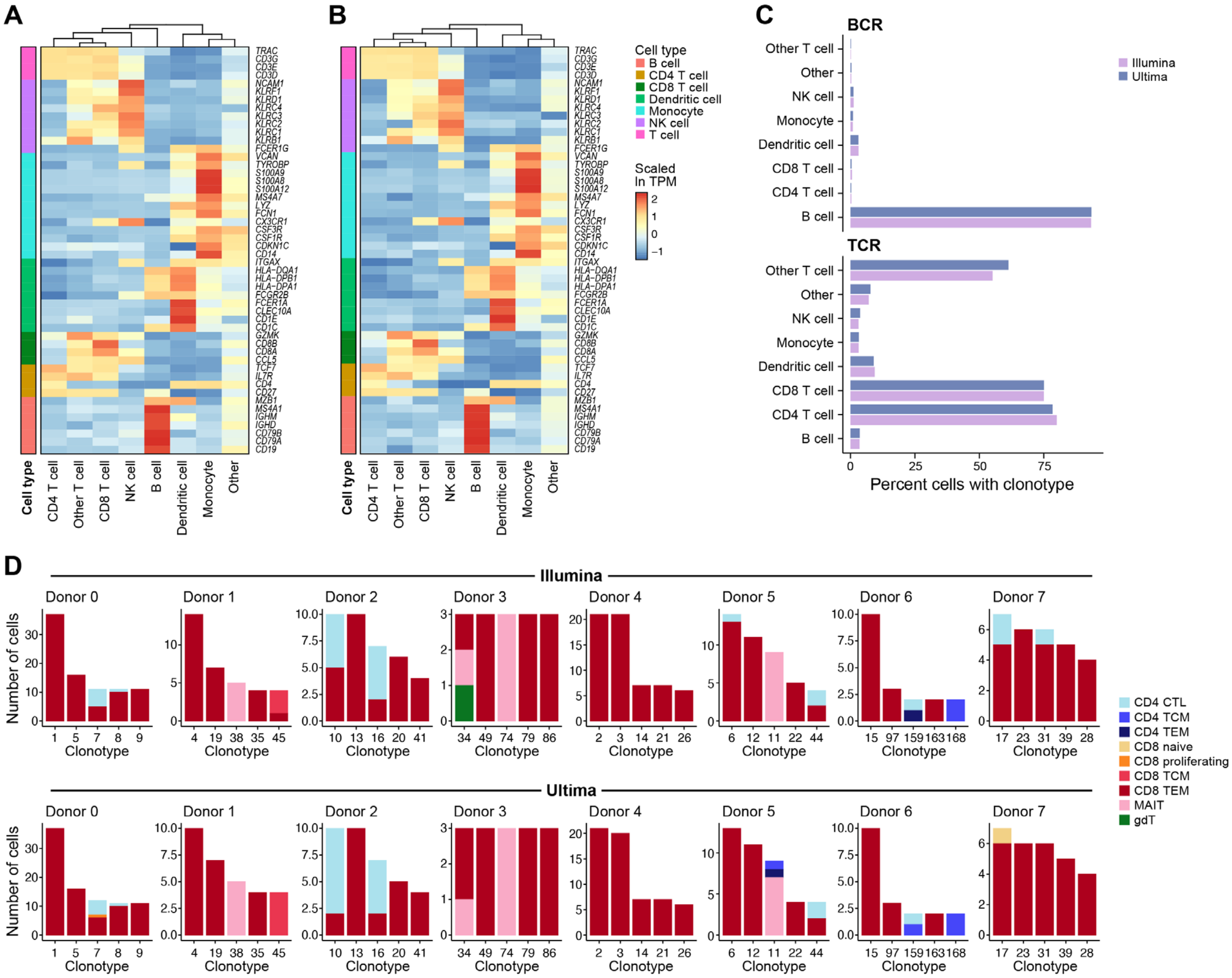

**Extended Data Fig. 6.**

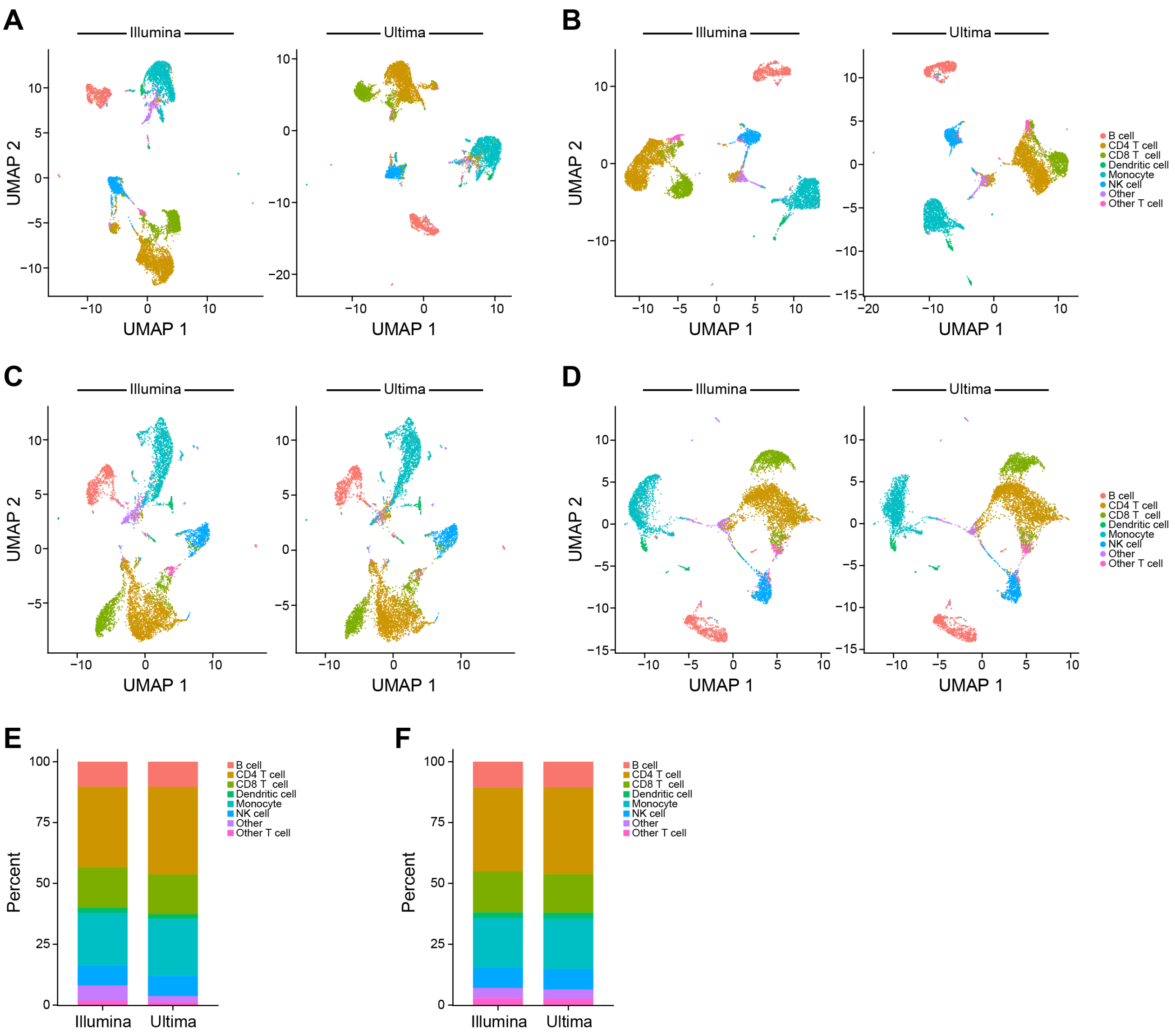


**Extended Data Fig. 7.**

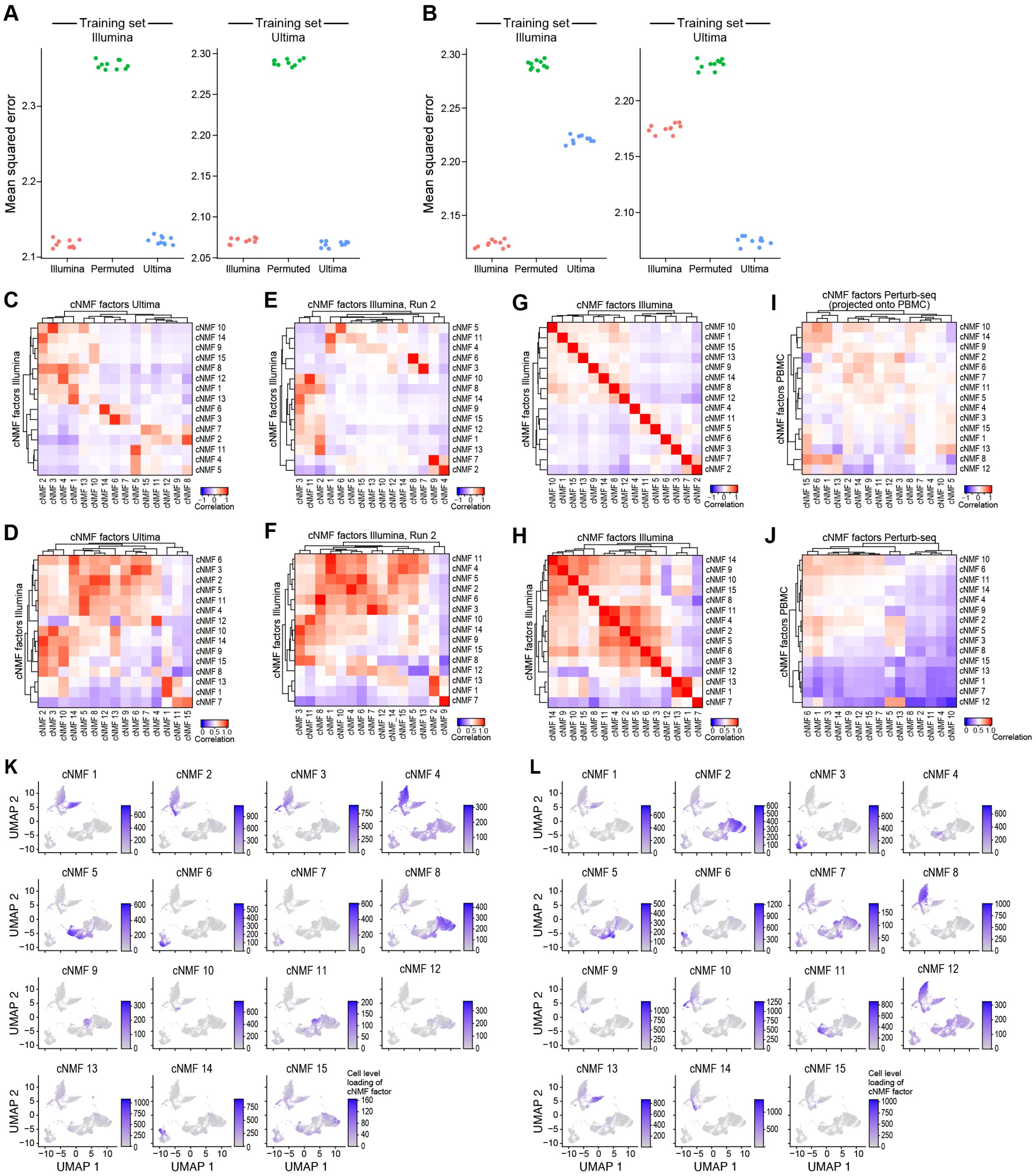


### Extended Data Fig. 8.

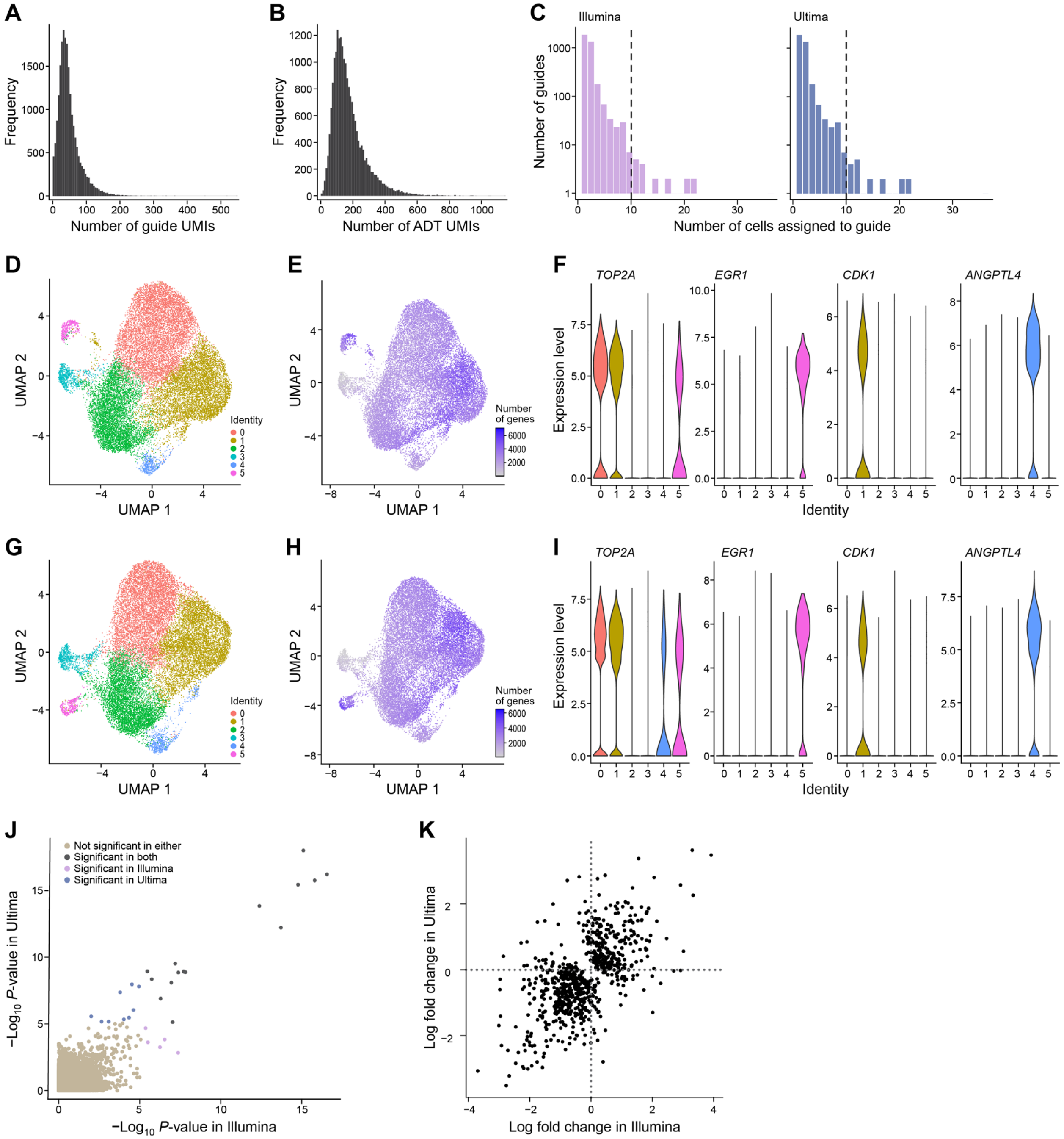


**Extended Data Fig. 9.**
